# Temporal resolution of spike coding in feedforward networks with signal convergence and divergence

**DOI:** 10.1101/2024.07.08.602598

**Authors:** Zach Mobille, Usama Bin Sikandar, Simon Sponberg, Hannah Choi

## Abstract

Convergent and divergent structures in the networks that make up biological brains are found across many species and brain regions at various spatial scales. Neurons in these networks fire action potentials, or “spikes”, whose precise timing is becoming increasingly appreciated as large sources of information about both sensory input and motor output. While previous theories on coding in convergent and divergent networks have largely neglected the role of precise spike timing, our model and analyses place this aspect at the forefront. For a suite of stimuli with different timescales, we demonstrate that structural bottlenecks– small groups of neurons post-synaptic to network convergence – have a stronger preference for spike timing codes than expansion layers created by structural divergence. Additionally, we found that a simple network model based on convergence and divergence ratios of a hawkmoth (*Manduca sexta*) nervous system can reproduce the relative contribution of spike timing information in its motor output, providing testable predictions on optimal temporal resolutions of spike coding across the moth sensory-motor pathway at both the single-neuron and population levels. Our simulations and analyses suggest a relationship between the level of convergent/divergent structure present in a feedforward network and the loss of stimulus information encoded by its population spike trains as their temporal resolution decreases, which could be tested experimentally across diverse neural systems in future studies. We further show that this relationship can be generalized across different spike-generating models and measures of coding capacity, implying a potentially fundamental link between network structure and coding strategy using spikes.

**Author summary:** Within the complex anatomy of the brain, there are certain structures that appear more often than expected. One example of this is when large populations of neurons connect to much smaller populations, and vice versa. We refer to these structural patterns as network convergence and divergence; they are observed in systems like the cerebellum, insect olfactory networks, visuomotor pathways, and the early visual system of mammals. Despite the ubiquity of this connectivity pattern, we are only beginning to understand its functional implications from a computational point of view. Here, we construct and analyze mathematical models of spiking neural networks to understand how convergent and divergent structure shapes the way that information is represented in each part of the network, as a function of the temporal resolution of population spiking activity. We then developed a simple feedforward network model of the visuomotor pathway of a moth, with similar convergent/divergent network structure, and reproduce a similar proportion of spike timing to spike count information as observed experimentally. Our results form predictions about spike coding in populations previously unobserved in experiment.

## Introduction

The neural systems of animals comprise networks with highly non-random topological structure [1–6]. The relationship between computation and connectivity in neural networks is multi-faceted and depends on our definition of these terms [7–9], but often it can be fruitful to focus on computation in networks with connectivity patterns that are observed more often in biological systems than would be expected in a totally random model network [4, 10–12]. One particular structural motif that is common in many areas of the nervous system involves populations of neurons synapsing with other populations of a much different size. When a large population of neurons synapses with a much smaller population, it may be called a “convergent” pathway. If a small population synapses with a much larger population, we call this structure “divergent.” Structural convergence and divergence are observed in a wide range of neural systems across species, including the mammalian early visual system [13–16], mammalian cerebellum-like structures [17, 18] and the insect olfactory system [19]. A notable example is the divergence from 200 million mossy fibers to 50 billion granule cells and then convergence to 15 million Purkinje cells in the human cerebellum – mostly a feedforward network. [20, 21].

Despite their ubiquity, convergent/divergent structures are only beginning to be understood from a functional point of view. Previous work has shown that network convergence synergizes with nonlinear activation functions to boost information coding [13]. Other studies have focused explicitly on networks with bottlenecks (small groups of neurons pre- and post-synaptic to much larger groups of neurons on both sides), demonstrating that modular connectivity increases their information transfer in classification tasks [22], and that they increase dimension while reducing noise in the expansion layer post-synaptic to them [17]. While highlighting the computational significance of structural convergence and divergence, the network or neuron models used in these studies were non-spiking, neglecting the biologically-relevant role of precisely-timed action potentials. Instead of directly testing how feedforward convergence and divergence shape coding strategies of spiking neurons, a recent study [23] examined the relationship between temporal coding of spiking population and its size. In this work, time-dependent stimuli were decoded from uncoupled spiking neuron populations of varying size. It was found that signal reconstruction error drops linearly with population size when decoded from precisely timed spikes, but sublinearly when decoded from imprecisely timed spikes. Although this work reveals an interesting relationship between spike coding and population size, it is still unclear how convergent and divergent network structures directly shape the importance of spike timing in information processing.

The demand for understanding the implications of convergent and divergent structure for the information coding of a spiking neural network is especially high in light of growing experimental evidence showing that spike timing can encode significantly more information about inputs [24] and outputs [25] than spike count. At the sensory input level, millisecond-level variations in spike timings encode significant proprioceptive [26], visual [27], auditory [28], olfactory [29] and tactile [30] information. We know now that similar levels of precision exist in the peripheral motor system. Millisecond-precise spike timings encode a significant (and sometimes greater than spike count) amount of information across human movement [31], muscle coordination [25], songbird acoustic structures [32] and respiration [33], insect flight control, turning maneuvers [25, 34], and escape behaviors [35]. Thus, in the sensory and motor peripheries of the nervous system, the importance of precise spike timing has been well established.

The role of spike timing is not as well understood in the intermediate stages of processing between sensory and motor populations, which, in the context of vertebrate and invertebrate visuomotor pathways, involve several cascades of structural convergence and divergence from the early visual system to cortex [15, 16] and eventually through the cerebellum [18, 20, 21] to the spinal cord and commanding muscles. A classic modeling study suggests that the cortex, a large population of neurons post-synaptic to structural divergence, is more likely to use a population spike count code due to high variability in inter-spike intervals [36]. Another work argues based on energy expenditure that rate/count coding can only explain around 15% of the activity in primary visual cortex [37], suggesting that other coding strategies must explain the rest [38]. From a purely quantitative perspective, single-neuron count codes are slow and information-poor, but robust to noise [39]. The activity of large populations of neurons following a structural divergence comprises a high-dimensional space and may therefore benefit from a collective count code due to the noise reduction. On the other hand, spike timing codes are fast, efficient, and information-rich, but potentially sensitive to noise [40]. It is, therefore, possible that bottleneck populations of neurons post-synaptic to structural convergences may be good candidates for a temporal code, since this would allow them to encode a similar amount of information as the larger pre-synaptic layer but with a smaller number of neurons. Indeed, experiments testing white noise optogenetic stimuli in the cortex of mice have shown that temporal precision of spiking increases in the inter-neurons post-synaptic to a structural convergence compared to the pyramidal neurons pre-synaptic to them [41]. However, this is just one example, and our understanding of the information processing between sensory input and motor output would improve if a relationship between population spike coding and convergent/divergent structure was also explored theoretically.

Thus, we aim to study this relationship systematically in spiking neural network models. Our primary hypothesis is that temporal coding is more beneficial in bottleneck populations post-synaptic to a structural convergence than it is in an expansion layer. While expansion layers have a surplus of neurons and may represent stimuli equally well with a coarse count code, bottlenecks have fewer neurons available to encode signals. Therefore, bottlenecks may preserve information by preferring temporal expressions of signal representations. To test this hypothesis, we train feedforward spiking neural networks to autoencode a time-dependent stimulus [38] and perform decoding analyses [42] on the population spike trains binned at various resolutions. First, we study 3-layered feedforward networks with varying levels of convergence and divergence to establish a relationship between structure and spike coding. Next, we develop a 5-layered model resembling the patterns of expansion and contraction in a hawkmoth visuomotor pathway, whose output is known to use a spike timing code during hover-feeding and target tracking [25]. We test if our model, although lacking many biological details present in the moth, can recapitulate a similar relative proportion of information in spike timing and spike rate coding as observed in experiment. To confirm that our results are not an exception due to the specific spiking model or decoder we chose, we also test the robustness of the results using other models and measures.

## Results

A graphical summary of our approach is shown in fig. 1. We first train a feedforward spiking neural network with a given structure to autoencode a time-dependent stimulus *s*(*t*) (left of fig. 1A), and then decode it using a recurrent neural network into its reconstruction *ŝ*(*t*) (bottom right of fig. 1A). To test how the encoding changes as we increase the temporal resolution of the spike trains, we use a decoding analysis in which we process each layer’s spikes over a sliding window of width *T* = 50 ms which are further binned at resolution Δ*t*. The choice of 50 ms for the duration of the response window was motivated by the wingstroke period of the hawkmoth *Manduca sexta*, and it is also consistent with previous neural decoding studies [42]. The binned spikes *R*(*t*; Δ*t*) are then fed to a decoder (the recurrent neural network) that treats the binned spikes within the larger response window as a sequence of hidden states within its own dynamics. The decoder estimates the stimulus presented to the input layer of the network with a reconstruction *ŝ*, based on the binned population spiking of the layer of interest. We then quantify the relationship between response *R*(*t*; Δ*t*) and stimulus *s*(*t*) by computing both the decoding accuracy *R*^2^ (coefficient of determination) and the mutual information *I*_*m*_ between true stimulus *s* and decoded stimulus *ŝ* for various Δ*t*. These measures approximate the true information carried at the population level and are computed across a range of Δ*t* to establish the temporal resolution of the optimal coding strategy, referred to here as the “information curves”.

**Fig 1.**
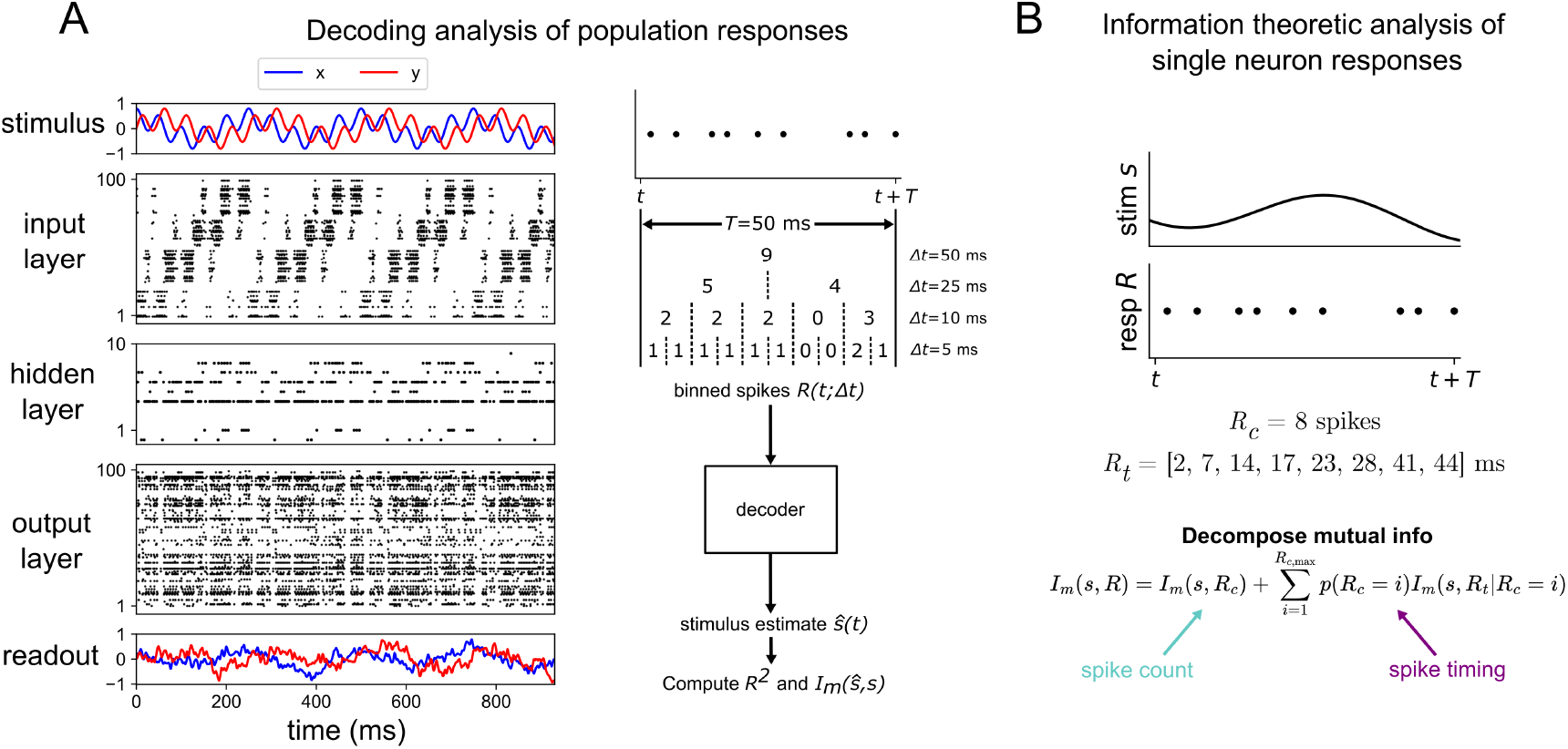
Schematic of model and analysis methods. (A) Raster plot of 3-layer network model trained to a 4 Hz + 20 Hz sum of sines stimulus. Red and blue indicate the *x*- and *y*-components for both the stimulus and readout (left). Depiction of the procedure used to process population spike trains before feeding them to the decoder to estimate the stimulus (right). *T* = 50 ms is the width of the sliding window used here and Δ*t* is the bin size (B) Sketch of the information theoretic method used to validate the 5-layer network model against previous results from hawkmoth data. A window of duration *T* = 50 ms is used in this analysis. The variable *R*_*c*_ denotes the spike count response and *R*_*t*_ denotes the spike timing response.

We also perform an information theoretic analysis at the single-neuron level, based on past work [25, 33, 43]. The strength of this method is that it quantifies the amount of spike count and spike timing information without confounding the two variables. Note that the binning method used in the population analysis considers spike counts over increasing levels of time resolution and therefore does not isolate spike-timing code completely from spike-count code. The strength of this method is that it considers all neurons in the layer and thus quantifies its population coding strategy, not just single-neuron coding. Computing mutual information at the single-cell resolution allows us to compare our model’s results with previously obtained experimental results at the single neuron level. For a detailed explanation of our model and analysis, see Methods.

### Three-layer network

We first focus on a feedforward network of 3 layers, systematically varying the number of neurons in the middle (hidden) layer *N*_*h*_ while keeping the number of neurons in the input and output layer fixed at *N*_in_ = *N*_out_ = 100. By doing this, we simultaneously tune the level of structural divergence and convergence. The network model consists of leaky integrate-and-fire neurons with both excitatory and inhibitory synapses. The parameters of the network, including synaptic weights and membrane time constants, are optimized to minimize the following loss function

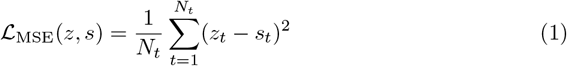

where *N*_*t*_ is the total number of time points, *s*_*t*_ is the true stimulus at time *t*, and *z* is a readout from the output layer of the form

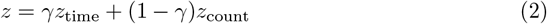

where *z*_time_ is a readout based on the spike timings of the output layer and *z*_count_ is a readout based on the spike counts of the output layer. The quantity γ is a hyperparameter that we set to 0.5, so as to equally weigh the readouts based on spike count and spike timing, thus not biasing our results (for more details, see Methods).

After training the network, we decode the stimulus from each layer by using the population spikes binned at various time resolutions Δ*t* using two types of recurrent neural networks (see decoding analysis methods). The association between the true stimulus *s* and decoded stimulus *ŝ* is estimated using various measures, including the mutual information *I*_*m*_(*s, ŝ*). Since the decoded stimulus is a function of the response (i.e. *ŝ* = *f* (*r*)), the data-processing inequality states that *I*_*m*_(*s, ŝ*) ≤ *I*_*m*_(*s, r*). Thus, when we quantify how associated *s* and *ŝ* are, we are computing a lower bound on the true association between stimulus *s* and response *r*. Note that the estimated stimulus *ŝ* from each layer and the network readout *z* are separate quantities: the former is constructed by binning population spike trains at various Δ*t*’s and feeding them to the decoder while the latter is purely a mechanism by which we train the network, thus increasing the information in the deeper layers before performing the decoding analysis which forms the *ŝ*’s.

For a variety of stimuli, we demonstrate how this information changes in each layer as a function of the network structure and timescale of spike counting Δ*t*. When Δ*t* is equal to the duration of the response window *T*, the input to our decoder is a vector of spike counts across each neuron. When Δ*t* = 1 ms (1 ms is the time step of our simulations), the input to the decoder is matrix of 1’s and 0’s indicating when spikes occurred at each time step, across all neurons in the population. Due to the loss in dimensionality of the neural representation implied by network convergence, we hypothesize that a temporal code (high information at small Δ*t* but low information at high Δ*t*) will be especially beneficial in bottlenecks. Conversely, large populations post-synaptic to network divergence should have less to gain from temporal codes, since they have high-dimensional representations even with a count or rate code (high information across all Δ*t*’s).

We first sought to test deterministic stimuli with fixed and well-defined frequency content, opting for sinusoidal stimuli of various frequency. We start with analyses of the optimal temporal resolution of codes at the *output layer*. In fig. 2, the information in the output layer has a steeper decline with growing Δ*t* in the case of the expansion hidden layer structure, as opposed to the bottleneck hidden layer structure, especially at higher stimulus frequencies. This is shown for a wide range of stimulus frequencies *f*_high_ in fig. 2C, where the slope of the information curves is plotted as a function of *f*_high_. There is a general decrease in the slopes with increasing stimulus frequency for both bottleneck and expansion networks, owing to progressively better encoding of faster stimuli by spikes binned at higher temporal resolution. Additionally, for all frequencies *f*_high_ *>* 20 Hz tested, the slope distributions are significantly lower in the expansion hidden layer structure (where signals converge onto the output layer) than the bottleneck hidden layer structure (signals diverge onto output layer). This demonstrates that structural convergence is associated with timing codes whereas structural divergence is associated with count codes. To ensure that this result did not depend on our specific choices, we tested different decoders and spiking neuron models in supplementary figures S1 and S2 and found the same result.

**Fig 2.**
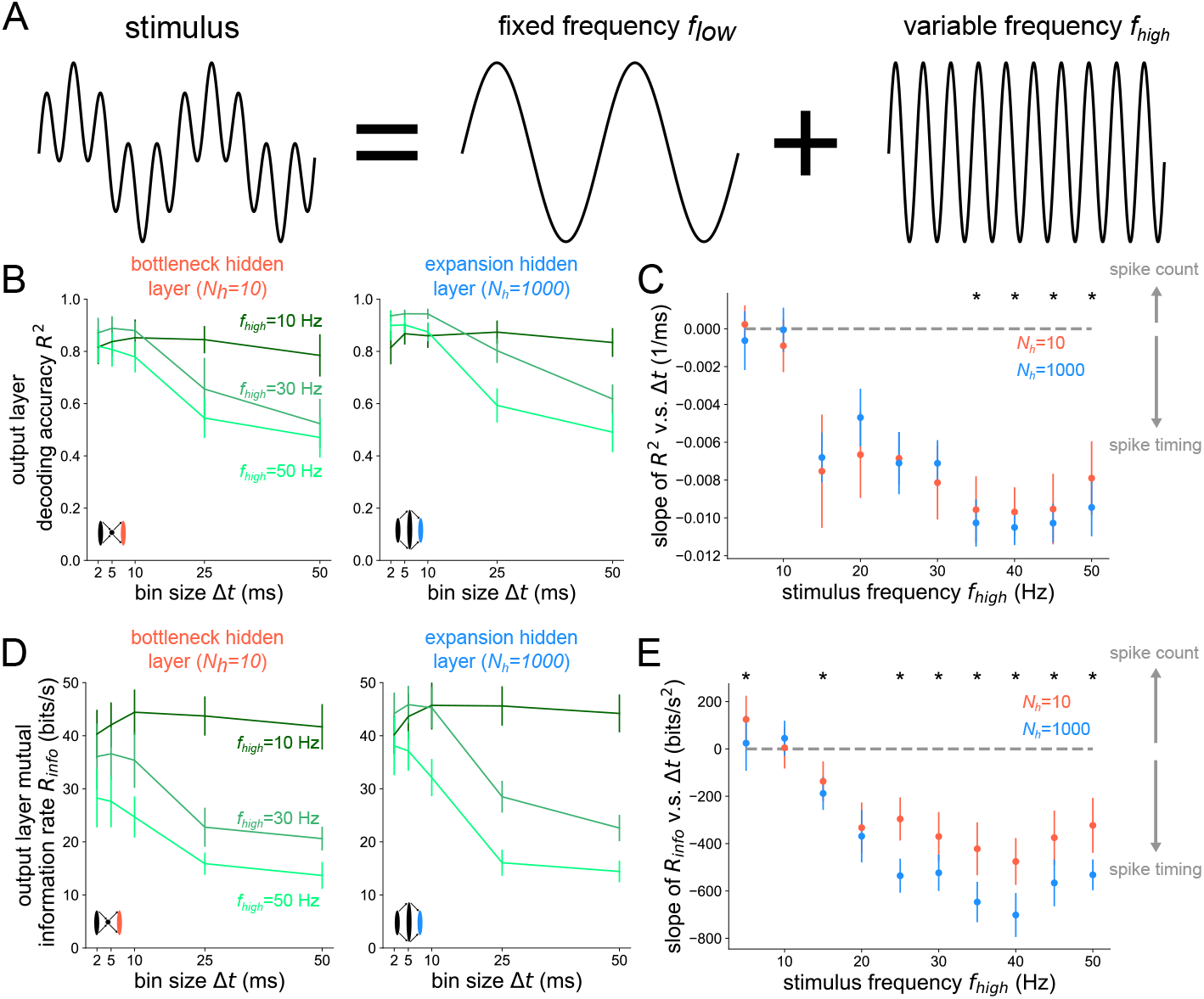
Structural convergence to the output layer promotes timing codes across stimulus frequencies. (A) Stimuli are sums of sines with fixed frequency component *f*_low_ = 4 Hz and variable component *f*_high_ (B) Decoding accuracy based on output layer spikes binned at time resolution Δ*t*. The left plot shows the result for *N*_*h*_ = 10 hidden neurons and the right shows it for *N*_*h*_ = 1000 hidden neurons. (C) Slope of *R*^2^ v.s. Δ*t* curves as a function of the high frequency stimulus component *f*_high_. Asterisks denote where a one-sided Wilcoxon rank-sum test is significant at *p* < 0.05. (D) Mutual information rate *R*_info_ = *I*_*m*_(*s, ŝ*)*/T* based on the output layer spikes binned at time resolution Δ*t*. (E) Slope of *R*_info_ v.s. Δ*t* curves as a function of *f*_high_, the high frequency stimulus component. Asterisks denote where a one-sided rank-sum test is significant at *p* < 0.05. Error bars represent distributions of the results over 10 independent network simulations.

For the same networks tested in fig. 2, we also performed a decoding analysis on the *hidden layer* for the case when *f*_low_ = 4 Hz and *f*_high_ = 20 Hz in fig. 3. As a visual representation of how more precise temporal codes are associated with bottleneck populations of neurons, stimulus reconstructions are shown for *N*_h_ = 10 and *N*_h_ = 1000 in fig. 3A from spike trains binned at Δ*t* = 5 ms and Δ*t* = 50 ms. In the case of an expansion hidden layer *N*_h_ = 1000, there is little difference between Δ*t* = 5 ms and Δ*t* = 50 ms; the drop in decoding accuracy when going from a more precise temporal code Δ*t* = 5 ms to a less precise code Δ*t* = 50 ms is only Δ*R*^2^ = 0.078 (see right side of fig. 3B). However, when decoding from the hidden layer of the bottleneck network *N*_h_ = 10, there is a large drop in decoding accuracy when going from a more precise code (Δ*t* = 5 ms) to a less precise code (Δ*t* = 50 ms). From fig. 3A, it is clear that the drop in accuracy comes from the fact that the *N*_h_ = 10, Δ*t* = 50 ms reconstruction misses the faster 20 Hz frequency component while the other reconstructions do not. By having a higher dimensional representation of the input, the *N*_h_ = 1000 expansion layer can still encode these higher frequency components even with a less precise code, binned over a time window equal to the period of the faster stimulus component. We again tested this result for an alternative spiking model, decoder, and association metric, finding the same general trend in supplementary figures S3, S4, S5, and S6.

**Fig 3.**
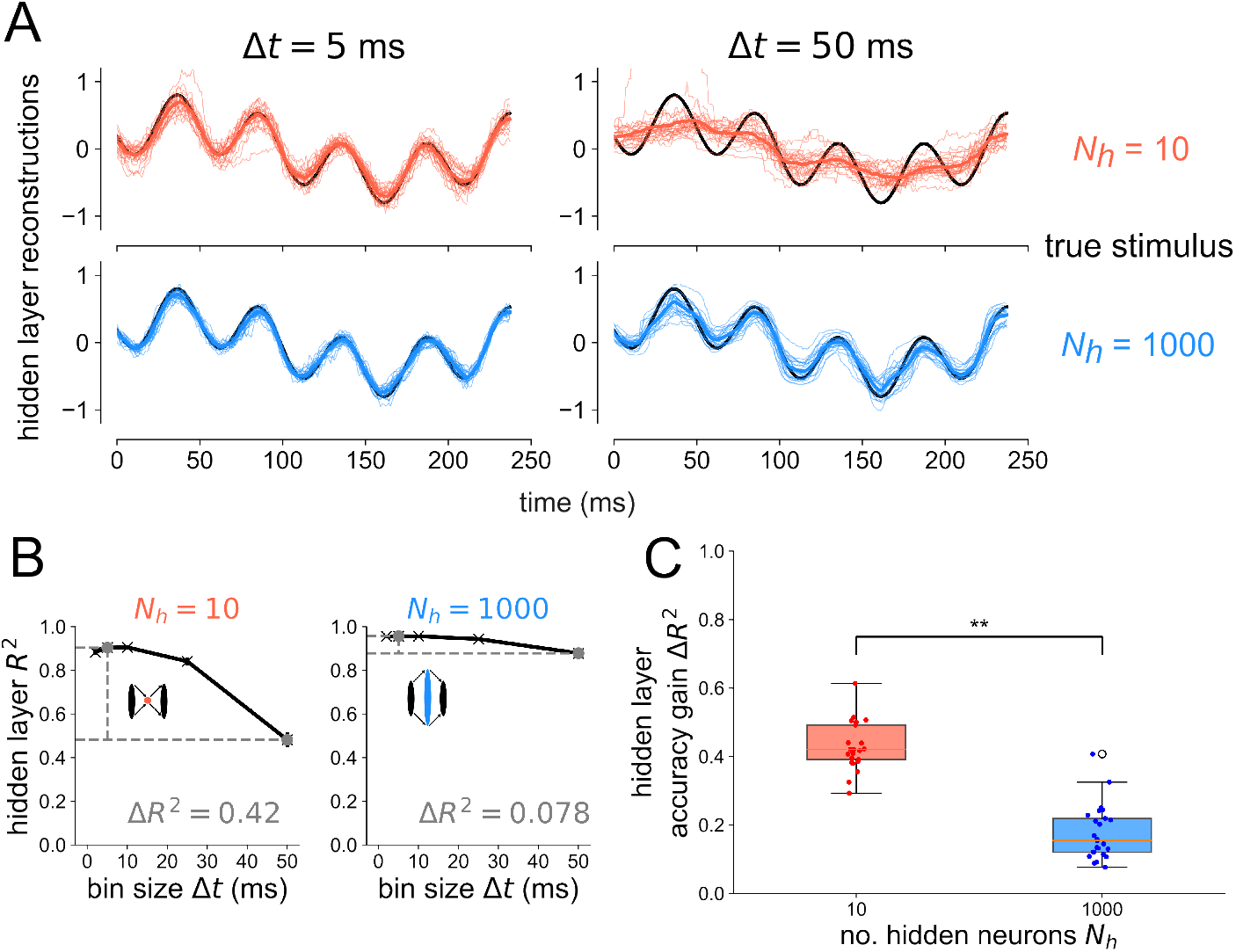
Bottlenecks have more to gain from temporal codes than expansion layers. (A) Example reconstructions from the hidden layer spikes binned at Δ*t* = 50 ms resolution for *N*_*h*_ = 10 (top) and *N*_*h*_ = 1000 (bottom). Thin traces show reconstructions from individual network seeds. Thick colored traces show means across all network seeds. (B) Decoding accuracy from the hidden layer spikes as a function of bin size Δ*t* for the bottleneck (left) and expansion (right) network. Gray points denote which bin sizes were used to compute accuracy gain Δ*R*^2^. Error bars denote standard errors of the mean over network seed distributions. (C) Accuracy gain of the temporal code over count code when reconstructing the stimulus based on spikes from the hidden layer, for bottleneck (red) and expansion (blue) networks. One-sided Wilcoxon rank-sum test *p* < 6 *×* 10^*−*10^. Results are shown for 25 independent network simulations.

To explicitly show that the higher-frequency component *f*_high_ = 20 Hz contributes to the drop in decoding accuracy for *N*_h_ = 10 at Δ*t* = 50 ms in fig. 3, we decode the low frequency component *f*_low_ = 4 Hz separately from the high frequency component *f*_high_ = 20 Hz in fig. 4 for all layers of the bottleneck and expansion networks. At the input layer (left), there is virtually no difference in the *R*^2^ v.s. Δ*t* plots between the bottleneck and expansion networks. When decoding from the hidden layer of either the bottleneck or expansion network, the decoding accuracy of the 4 Hz component remains constant for all Δ*t*’s. However, there is a large discrepancy between the bottleneck and expansion networks when decoding the 20 Hz component from the hidden layer: the bottleneck has a steep decrease in decoding accuracy with increasing Δ*t* while the expansion shows a much slower decrease in *R*^2^ with increasing Δ*t*. Furthermore, going from the hidden layer to the output layer steepens the 20 Hz curve for the network with an expansion hidden layer, but leaves the 20 Hz curve for the network with a bottleneck hidden layer virtually unchanged. These results support the conclusion that populations post-synaptic to a network convergence encode high-frequency stimulus information with spike codes of high temporal resolution. Populations post-synaptic to structural divergence maintain similar information curves as their pre-synaptic layer, indicating that either spike count or spike timing codes are feasible for divergent populations.

**Fig 4.**
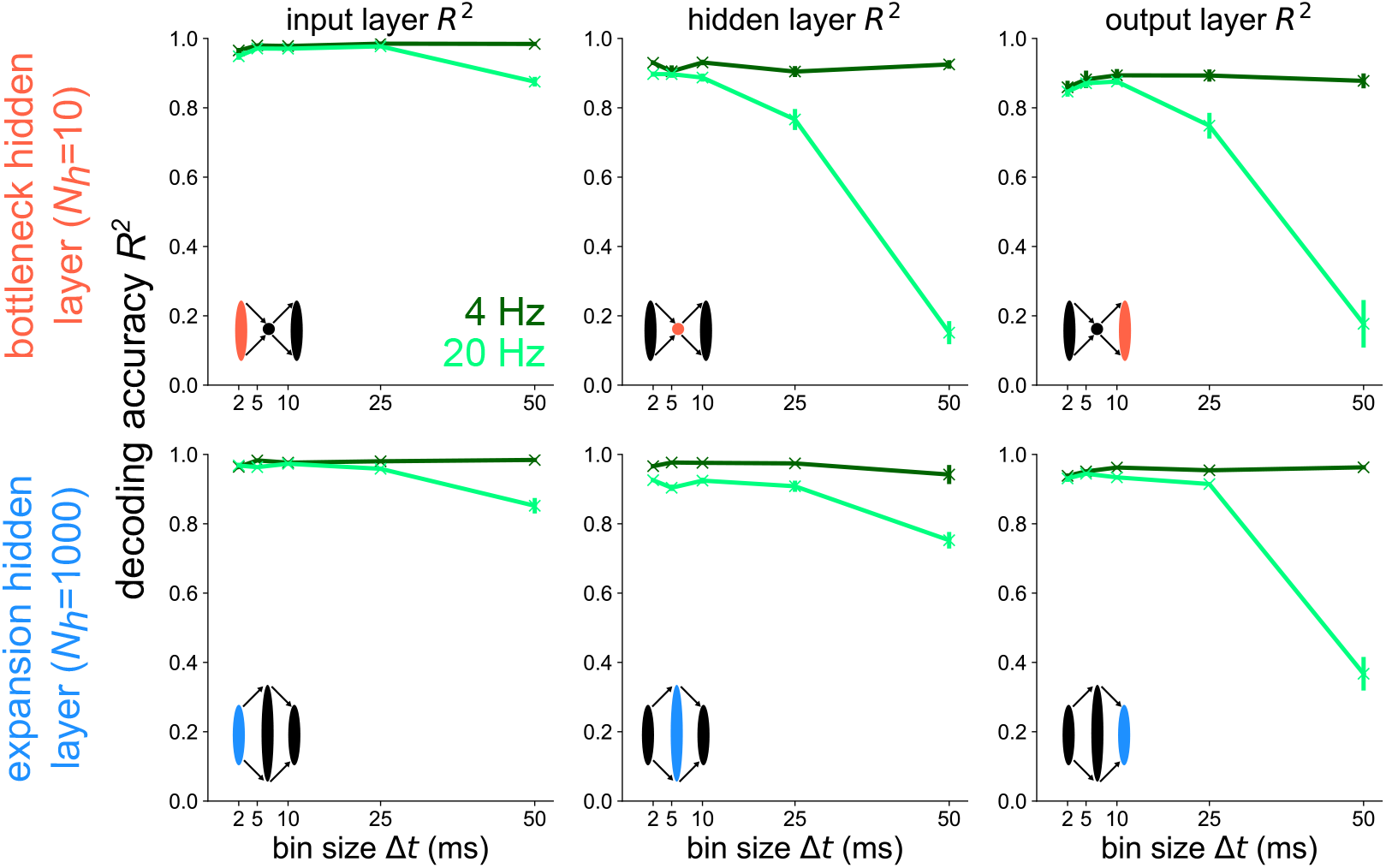
Temporal codes capture high-frequency stimulus components stronger in layers following structural convergence. Decoding accuracy versus bin size for each layer of the bottleneck (top) and expansion (bottom) networks receiving a 4 Hz + 20 Hz sum of sines stimulus. The 4 Hz (dark green) and 20 Hz (light green) components are decoded separately here. Error bars represent standard errors of the mean over 10 independent network seeds.

In the previous results, all stimuli used were sums of 2 sines. In fig. 5, we show accuracy gains in the hidden and output layer for four different stimuli. For a slow (5 Hz), continuous single sine stimulus (top), there is little to be gained from a more precise temporal code. For the other stimuli shown, which all include some sort of faster time scale or unpredictability, the hidden layer has a higher accuracy gain in a bottleneck network than a uniform (*N*_h_ = 100) or expansion (*N*_h_ = 1000) network. For the white noise and binary stimuli, the output layer has significantly higher accuracy gains in the expansion-hidden-layer network (*N*_*h*_ = 1000) than in the networks without structural convergence onto the output layer. Together, these results demonstrate that structural convergence promotes temporal coding in networks responding to stimuli with fast timescales or unpredictability. For slow stimuli without fast jumps, there is little, if anything, to be gained from a temporal code for all network structures tested.

**Fig 5.**
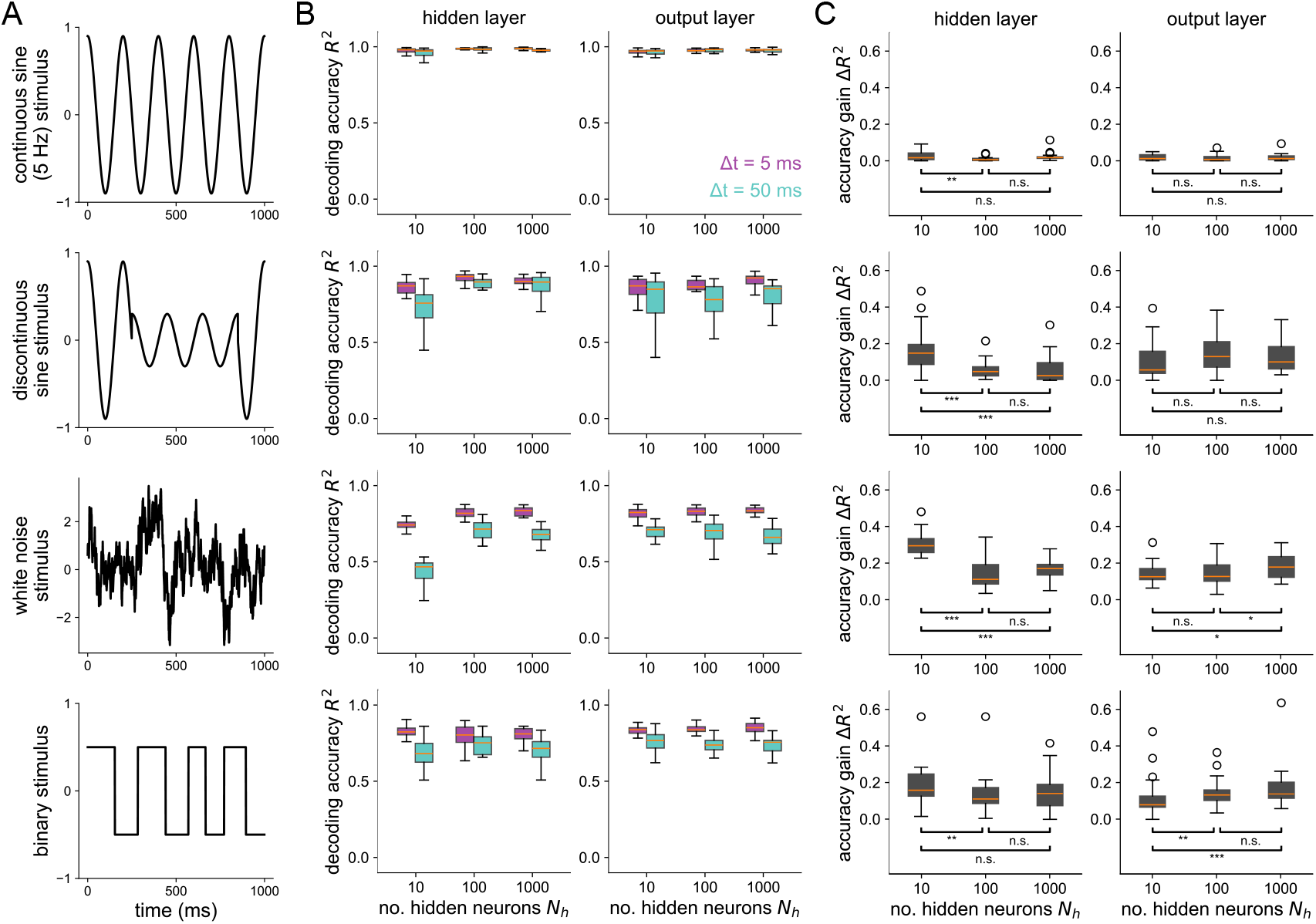
Stimulus-dependence of spike coding as shaped by convergent/divergent structure. (A) Each row shows the stimulus used for the corresponding plots on the right. (B) Decoding accuracy v.s. the number of hidden neurons at Δ*t* = 5 ms and Δ*t* = 50 ms for the hidden layer (left) and output layer (right). (C) Accuracy gain (*R*^2^ at Δ*t* = 5 ms minus *R*^2^ at Δ*t* = 50 ms) v.s number of hidden neurons. Asterisks denote where a one-sided Wilcoxon rank sums test is significant (* for *p* < 0.05, ** for *p* < 0.01, and *** for *p* < 0.001). All boxplots represent distributions of the results over 25 independent network simulations.

### Five-layer network model of the moth visuomotor pathway

Now that a relationship between optimal coding strategy with spikes and convergent/divergent structure has been established in a simple 3-layer model, we next test our model-based conclusion on this relationship in a specific biological model of a convergent/divergent neural network found in nature. Specifically, we focus on the visuomotor pathway of the hawkmoth *Manduca sexta* for its convergent/divergent architecture along the signal pathway and relative behavioral simplicity during flower tracking [44]. The output of this system primarily consists of only 10 muscles that control wing motion, each acting effectively as a single motor unit or output channel. This compact set of muscles, recorded with spike-level resolution, encode the majority of the information about motor output in their spike timing [25, 45].

The output layer of the hawkmoth visumotor pathway provides a nearly complete motor program for behavior allowing for the near perfect (*>*99%) reconstruction of behavioral output states [46] and between 85% and 95% reconstruction on the continuous 6 degree of freedom (DoF) body forces and torques [47, 48]. The input layer corresponds to the visual system, which we have here simplified as a group of 48 motion-sensitive neurons [24] separated into two subpopulations, each tuned to a direction along a line. Intermediate layers of the moth’s visuomotor pathway include the brain, the neck connective, and the thoracic motor circuits that drive wing muscles.

Structurally speaking, each of these populations corresponds respectively to an expansion (from 10^5^ to 10^6^ neurons), a bottleneck (big convergence from 10^6^ to 10^3^ neurons), and another expansion (from 10^3^ to 10^4^ neurons) before finally converging (from 10^4^ to 10^1^ neurons) at the output layer. For a schematic diagram of the moth’s visuomotor pathway and our corresponding model network, see fig. 6. The size of each neural population in the model network was chosen to preserve the relative order of magnitude of divergence and convergence observed in the moth, within computational capacity. This is a very coarse representation of the real network. Of course many brain regions are not involved in the process of target tracking but the optic lobe and premotor regions capture a very large portion of the brain of moths and other insects [49–52].

**Fig 6.**
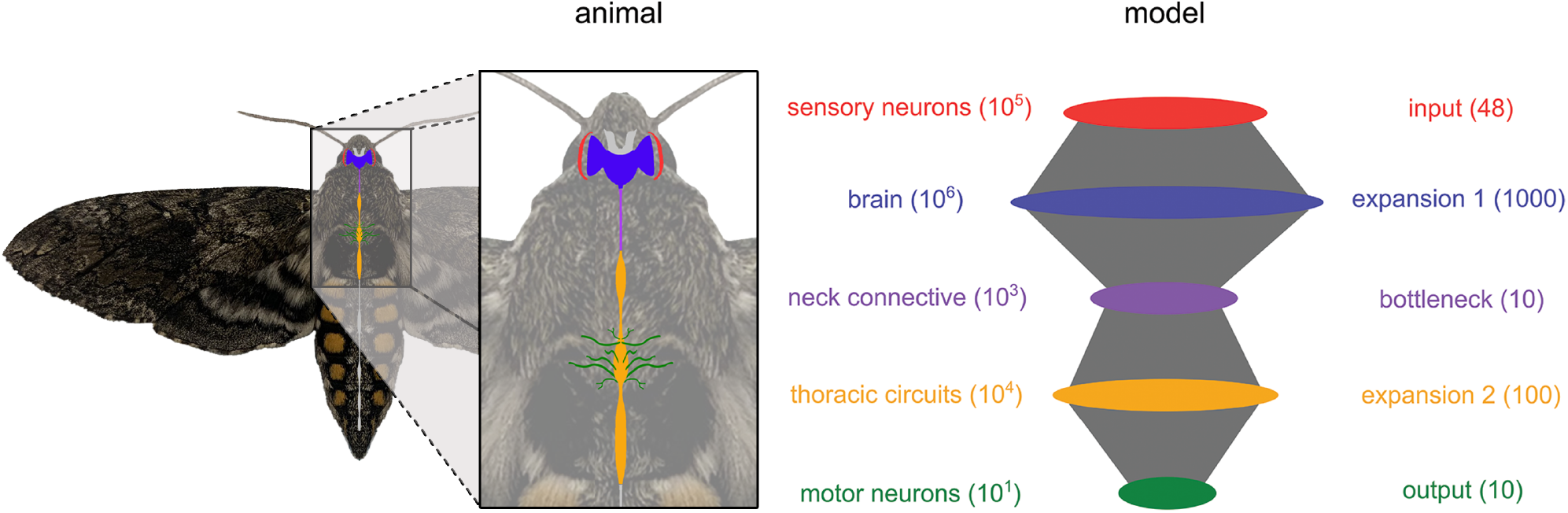
Experimental system and network model. Diagram of the central nervous system of the hawkmoth *Manduca sexta* and a schematic of the 5-layered spiking neural network developed here to model its visuomotor pathway. Numbers in parentheses denote the number of neurons in each population for the moth (orders of magnitude, left) and the model (exact, right).

Our first objective with the 5-layer network was to validate it against previous findings from the motor program of the hawkmoth. In particular, Putney et al [25] performed experiments where hawkmoths were shown a robotic flower oscillating horizontally at 1 Hz, a stimulus that they are naturally inclined to track when foraging. During the flower tracking, the 10 muscles coordinating their flight were recorded with spike timing resolution down to 0.1 ms. The authors found that a significant majority of the mutual information between the spiking activity of these muscles and the motor output (forces/torques generated during flight) was encoded by spike timing instead of spike count in each unit. In fact, spike timing encoded three times more information than spike count. Subsequent analysis showed that the precision of the spike timing code was of the order of 1 ms across all output units [45].

We re-analyzed their data from moth motor units to first confirm this result, shown in fig. 7C. Next, we trained our 5-layer network model on a 1 Hz stimulus that was used during the experiment and performed the same single-neuron information theoretic analysis for all layers of the model. Since there is no “motor output” from our model, we computed the mutual information between the stimulus and the response, which is analogous to the mutual information between motor output and response in a setting where the stimulus is being physically tracked. The results are shown in fig. 7. In particular, a large majority of the mutual information in the output layer of our model is encoded by spike timing (bottom of fig. 7C), just as found from the experimental data (top of fig. 7C). Furthermore, we show the single-neuron information rate averaged across all neurons within a layer in fig. 7B. The spike count information is low in all layers compared to the spike timing information. The single-neuron spike timing information starts low in the input layer, rises in the first expansion (E1) layer, falls in the bottleneck (B), rises slightly in the second expansion (E2) and again in the output layer. In the output layer, the spike timing information rate exceeds the spike count information rate by a much larger amount than it does in the input layer. Note that the information theoretic method used in this analysis is conservative in the sense that contributions from spike timing are only taken once those from spike count have been completely accounted for. Overall, our result lends evidence to the notion that convergent/divergent structure in the hawkmoth visuomotor pathway supports a transformation from the input layer where spike timing is less important to the output layer where spike timing provides an order of magnitude more information than spike count. Furthermore, when interpreted in light of the population decoding analysis showing perfect reconstruction across all Δ*t*’s in all layers (supplementary figure S7), the single-neuron analysis shown in fig. 7B indicates that there is a high amount of redundancy in the large expansion layers. We also confirmed that pairwise redundancy in the output layer of the model is mostly contained in spike timing, not spike count (supplemental figure S11), which was another key result of ref. [25] demonstrating that coordination in hawkmoth hovering is achieved through spike timing and not spike count.

**Fig 7.**
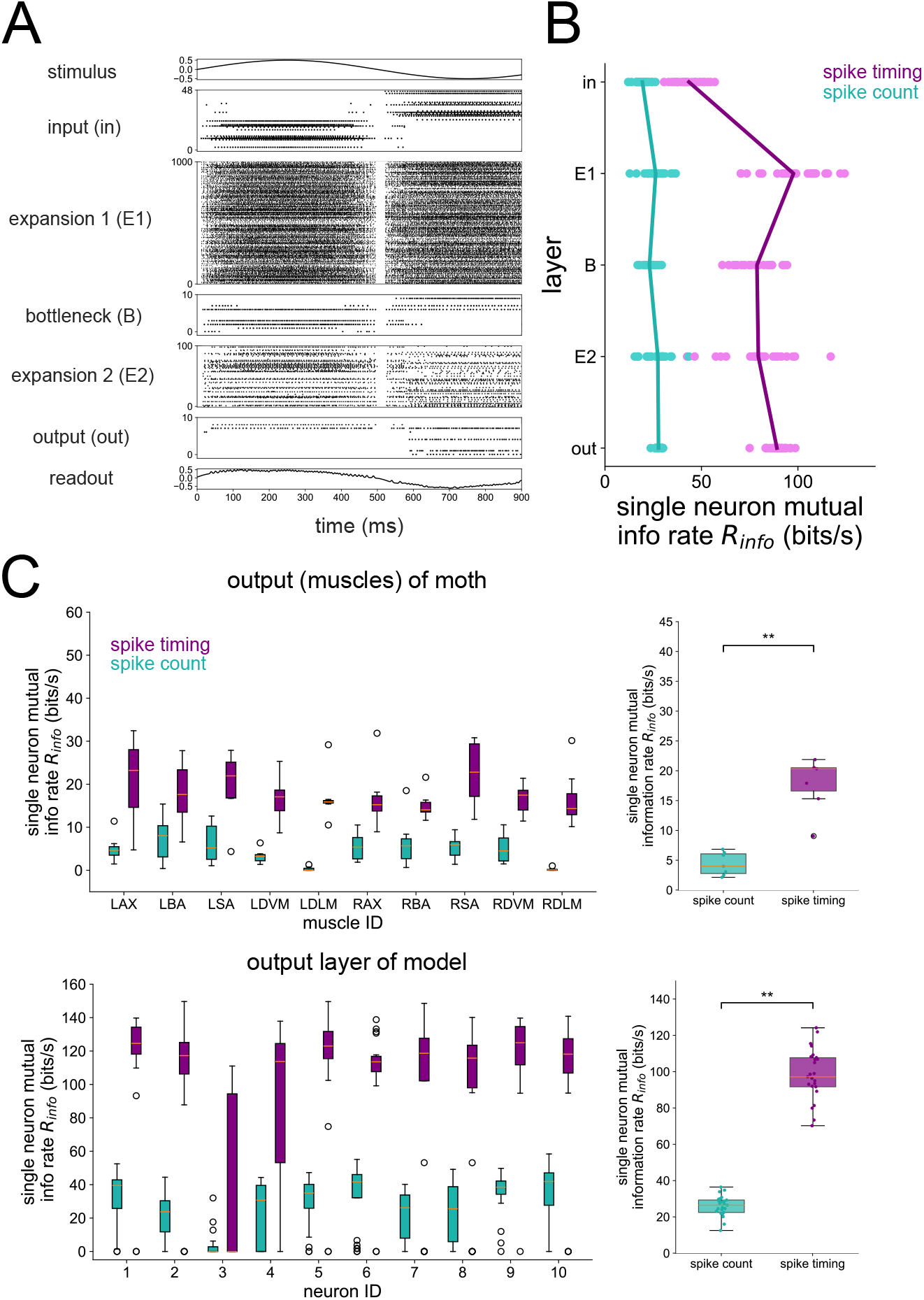
Single-neuron information during 1 Hz stimulus. (A) Raster plot of the 5-layer network model trained to a 1 Hz sinusoidal stimulus. (B) Single neuron information rate in each layer, decomposed into spike count and spike timing contributions. Each dot represents the result of a single network seed, averaged across all neurons in the layer. Lines connect the means of the distributions. (C) Mutual information in spike count and spike timing from the hawkmoth motor program (top) and the 10 neurons in the output layer of the model (bottom). The plots on the right show mutual information pooled from the output muscles (top) and output layer of the model (bottom). Asterisks denote where a one-sided Wilcoxon rank sums test is signifiant at *p* < 0.01. For the model, mutual info *I*_*m*_(*s, R*) is taken between stimulus and response; for the moth data, mutual info *I*_*m*_(*m, R*) is taken between motor output *m* and response. The single-neuron method depicted in fig. 1B and described in Methods was used here to compute mutual information, consistent with Ref. [25], which is where the moth data was originally published. For the moth muscle results, boxplots represent distributions over 7 individual moths. For the model results, boxplots represent distributions over 25 independent network simulations.

Since the 1 Hz sinusoid was decoded very well in all layers and at all time scales (see supplementary figure S7), we sought to investigate what coding strategy was optimal during a more complex and biologically-relevant stimulus. Specifically, we were interested in the idea that the bottleneck may filter the noise in some way. To answer this, we trained the 5-layer network on a noiseless 4 Hz + 20 Hz sum of sines stimulus. Its input was a version of the same stimulus but with white noise added. In each layer, we decoded the noiseless stimulus from population spikes binned at various Δ*t*’s, the results for which are shown in fig. 8. We found that both expansion layers have a broader range of Δ*t*’s than the smaller layers over which nearly perfect decoding accuracy is achieved. This was quantified by computing the slope of the best line fits to the *R*^2^ v.s. Δ*t* curves shown on the top of fig. 8B. The distributions of these slopes for each layer are shown in the bottom of fig. 8B, and also explicitly against layer in fig. 8C. A slope of zero means that there is no preference for spike count or spike timing. A negative slope indicates that there is a gain in information with a spike timing strategy over a spike count coding strategy. For the noisy sum of sines used here, all of the slopes (except for one outlier in the E1 layer) were negative. However, the slopes were more negative in the bottleneck and output layer, supporting the conclusion that structural convergence promotes temporal spike coding.

**Fig 8.**
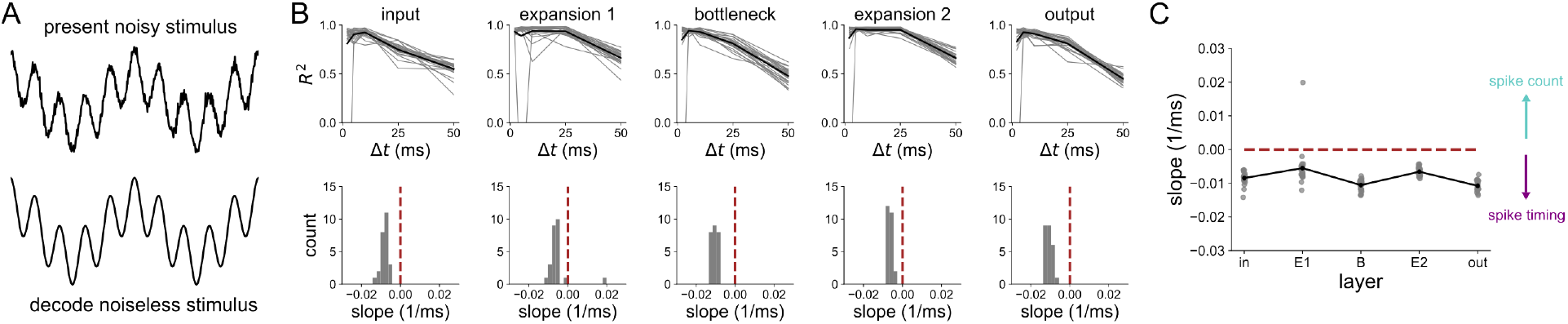
Decoding analysis of a noisy 4 Hz + 20 Hz stimulus. (A) The 5-layer network receives a sum of sines corrupted by noise, but is trained to encode the noiseless version at the output. The decoding is done with respect to the noiseless stimulus. (B) Decoding accuracy from spikes binned at resolution Δ*t*, in each layer of the 5-layer model. Each gray trace represents an individual network seed. Black traces are the means across all network seeds (top). Distribution of slopes of best line fits to the *R*^2^ v.s. Δ*t* curves (bottom). (C) Slope distributions versus layer. Results are shown for 25 independent network simulations.

## Discussion

We observe significant differences in information between spike count and spike timing representations as a function of convergent/divergent network structure. Although the stimulus reconstruction task is relatively low-dimensional, the fact that we are decoding from discrete spikes and not continuous rates makes this problem more difficult. Nonetheless, we notice differences in performance between spike count and spike timing representations, even for large layers. The 3-layer network results show that bottleneck populations of neurons post-synaptic to a structural convergence have more to gain from precise spike timing codes than expansion layers, so long as the stimulus being encoded has sufficiently fast dynamics. The simple 5-layer network model replicating the cascades of convergence and divergence in the moth sensory-motor pathway can reproduce the relative proportion of spike timing information previously measured at the single unit level from the spike resolved motor program of *Manduca sexta*. Notably, the amount by which spike timing information exceeds spike count information at the output layer is higher than that at the input layer. Even without the extensive recurrence and reafferent sensing observed in biological networks, our simple feedforward model replicates the experimental result at the hawkmoth motor output. This suggests that the feedforward signal compressions and expansions induced by the structural convergence and divergence can predict the relative information gain obtained by temporal coding, along the various stages of the hawkmoth visuomotor pathway. Our work goes beyond previous theoretical studies considering the effects of convergent and divergent structure on information processing [13, 17, 22] by establishing a relationship between this ubiquitous structural motif and the information encoded by spikes at various time resolutions in its constituent neurons.

A related but distinct concept to structural bottlenecks in biological neural networks is that of the information bottleneck: a variational method for extracting the most relevant information that a random variable *X* has about another random variable *Y* by finding an optimal compressed representation 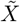 [53]. This method optimizes the tradeoff between prediction and compression and has been used to shed light on learning [54] and optimal architectures [55] in deep neural networks. While the vanilla information bottleneck method is agnostic to any particular mapping between 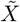 and *Y*, recent work has extended the idea by finding an 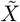 that is specific to the decoder being used for downstream prediction [56], thus improving generalization in artificial neural networks. A similar variant of the information bottleneck was applied to neural data from the cells of the retina, showing that predictive information about future visual inputs can be encoded and compressed by neurons post-synaptic to the retina [57]. Although the information bottleneck method is useful for understanding artificial neural networks [54–56] and neural data [57], its potential mapping to the discussion of structural bottlenecks in biological neural networks is unclear. Whereas information bottlenecks are optimal compressions in an abstract sense, the structural bottleneck studied here is a feature of networks widely observed in biology that we treat as a starting point and study its consequences for information-processing.

Network convergence and divergence are widespread in species and brain areas [14–16, 18, 19], but the implication of this structure for spike coding of time-dependent stimuli has not been well characterized. While the importance of spike timing at both sensory input [26–30] and motor output [31–35] is well established, its role in the intermediate processing stages resulting from structural convergence and divergence has been less clear [36, 37, 58, 59]. Our results demonstrate that bottlenecks would benefit greatly from a more temporally-resolved spike code, more so than in expansion layers which have a plethora of neurons to represent a signal with spike counts. This finding is relevant to a variety of systems where cascades of network convergence and divergence are present, including visuomotor pathways, cerebellum-like structures, the early visual system, and olfactory systems [13, 16–18]. Additionally, the nervous systems of segmented organisms contain neural ganglia which are often coupled by fewer fibers than they comprise, resulting in a convergent/divergent connectivity pattern [60].

Recent theoretical work with groups of uncoupled Poisson neurons is consistent with our finding that larger populations of neurons can encode time-dependent signals well with a count code, whereas small populations must use precisely-timed spikes to achieve the same decoding accuracy [23]. Here we show that this trend extends to feedforward networks with convergent/divergent structure where the number of neurons pre-synaptic to a given population shapes the coding strategy of that population, even when the size of that population is fixed (as in fig. 2). Additionally, while our decoder is a nonlinear function (a recurrent neural network) trained on discrete sequences of population spiking, the decoder used in Ref. [23] is a linear function of spikes convolved with an exponential filter of width 10 ms. Thus, our method accounts for sequences of population spike trains extended in time whereas the method used in Ref. [23] decodes a continuous signal from a continuous representation of a spike train over a short time. Our technique is more consistent with emerging definitions of spike timing codes in which longer sequences of spikes are critical for encoding information. [24, 25, 61, 62]. In order to obtain elegant analytical results, the authors of Ref. [23] assumed that their neural populations were not correlated with the signal being decoded, whereas the networks in our computational study explicitly encode the stimulus in the input layer and are trained to encode it at the output layer.

Whether our findings could be recapitulated in alternative learning models is an open question. Although artificial neural network (ANN) models can predict precise spike timing from biologically-relevant stimuli [48], they are unable to make predictions for the role of spike timing in intermediate layers since their units do not have a spiking mechanism. Indeed, this frontier is where past work has delivered mixed results [36, 37, 58] and was important for us to test. Other approaches like the “chronotron” [63] and “tempotron” [64] are single-neuron models that learn to classify inputs with distinct spike timing patterns. However, a training method such as this, although useful in other contexts, would bias the coding strategy toward spike timing, which is undesirable when interested in isolating the effect of network structure on coding in a study like ours. For this reason, we chose to train our network in a way that was agnostic to the coding strategy at the output (see eq. (2)), a notable strength of our approach.

There are many models of spike-generating mechanisms for the design of SNNs and these may promote different coding and network features. For example, the dynamics of resonant-and-fire neurons [65] and their generalizations [66] are selective for stimuli of certain frequencies. This could be especially important in the context of spike timing codes, since patterns of pre-synaptic spikes that resonate with the natural frequency of the post-synaptic neuron will more reliably result in that neuron firing [67]. This is in contrast to leaky-integrate-and-fire (LIF) neurons, which are most likely to spike when the input amplitude is high and the frequency is low. Although we tested models only within the LIF family, this particular model in its generalized form has been shown to reproduce a variety of experimentally measured neuronal spiking behaviors [68]. Thus, we expect that the general trends we observe in the two LIF models tested here will extend to other spike-generating mechanisms.

In summary, we found that convergent and divergent structure shapes the way in which populations of neurons encode high-frequency or less predictable dynamic stimulus information with precisely-timed spikes. Structural bottlenecks resulting from network convergence benefit much greater from precise spike timing than expansion layers coming from network divergence. A simple model recapitulates previous experimental findings at the motor output of the visuomotor pathway of the hawkmoth. While comprehensive experimental data across all layers of the hawkmoth visuomotor pathway is unavailable, our model further makes predictions about unobserved populations and untested stimuli, which could be confirmed experimentally in future studies. In particular, our single-neuron analyses predict high amounts of redundancy in the spike timing representation of simple visual stimuli by the populations comprising the brain and thoracic circuits of the hawkmoth. From our population decoding analyses, we predict that precise spike timing representations for accurate tracking of fast stimuli are needed in the bottlenecks of the hawkmoth visuomotor pathway (neck connective and motor neurons) but provide only marginal gains over spike count codes in the expansion layers (the brain and thoracic circuits). Overall, our work establishes a novel structure-function relationship in feedforward neural networks with signal convergence and divergence, elucidating how this structural motif prevalent across neural systems and species determines the optimal coding strategy with spikes.

## Methods

### Analytical support for spike timing and count codes

In this section, we present an analytical explanation for why network convergence promotes timing-based codes, demonstrating that count-based coding requires more neurons to achieve the same entropy upper bound as timing-based coding.

### Single-neuron example

Consider the example where the response window is of duration *T* ms and the refractory period of the neuron is *τ*_ref_ ms. The total number of bins to place spikes in would then be *n*_bins_ = *T*/*τ*_ref_. In the case of a spike count code, we may bin spikes at resolution Δ*t* = *T* ms. Including the outcome of 0 spikes, the total number of outcomes for the spike count code is |*S*_*c*_| = *n*_bins_ + 1. By assuming each of these outcomes is equally likely, the probability distribution becomes uniform, i.e. the probability of *i* spikes is *p*_*i*_ = 1/|*S*_*c*_| for *i* = 0, 1, …, *n*_bins_. Using this probability distribution with maximum entropy, we may calculate an upper bound on the true entropy of the spike count code. Let us refer to the true entropy of the spike count code as *H*_*c*_ and its upper bound as 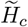. Then we have:

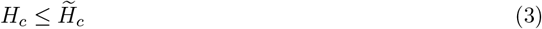

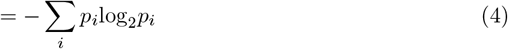

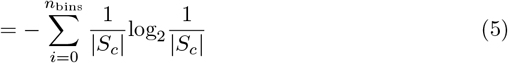

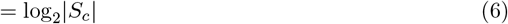

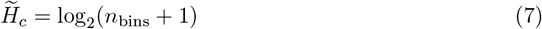

Similarly, we may bin spikes at resolution Δ*t* = *τ*_ref_ ms and list the possible spike timings as binary sequences. The total number of possible outcomes for the spike timing code is equal to the number of binary sequences of length *n*_bins_, which is given by |*S*_*t*_| = 2^*n*^bins. Again assuming a uniform distribution *p*_*i*_ = 1/|*S*_*t*_| for each outcome *i* = 1, …, |*S*_*t*_|, the upper bound on the entropy of the spike timing code is:

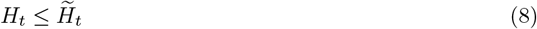

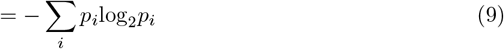

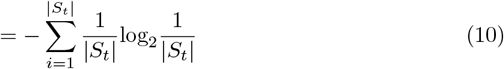

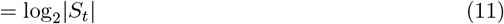

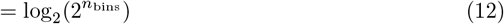

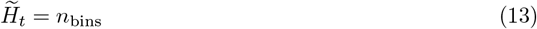

Therefore, the entropy upper bound for spike count code scales logarithmically with the duration of the response window whereas that for the spike timing code scales linearly. For the example when *T* = 15 ms and *τ*_ref_ = 5 ms, the number of bins is *n*_bins_ = *T/τ*_ref_ = 3 and the maximum entropy rate is 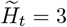 bits per 15 ms for the spike timing code and 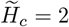 bits per 15 ms for the spike count code.

### Population of neurons

Let us now consider spike coding in a population of *N*_nrn_ neurons. In the case of a spike count code, each neuron in the population can fire anywhere between 0 and *n*_bins_ spikes. Therefore, the total number of outcomes is 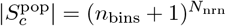. The upper bound of the population spike count code entropy is then:

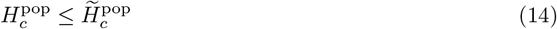

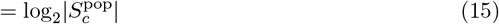

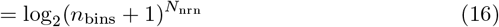

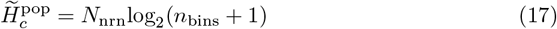

For the spike timing code, each neuron in the population can fire one of 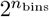 possible spike sequences. Thus, the total number of possible outcomes for the population spike timing code is 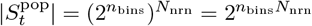. The upper bound of the entropy for the population spike timing code is:

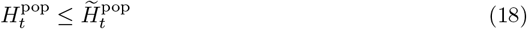

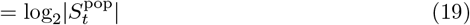

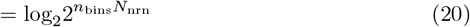

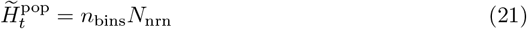

The entropy upper bounds for both the spike count and spike timing codes grow linearly with the number of neurons *N*_nrn_ but with different slopes. For the spike count code, the slope is log_2_(*n*_bins_ + 1). For the spike timing code, the slope is *n*_bins_. The slopes of the entropy upper bounds of both the spike timing and spike count codes are plotted as a function of *n*_bins_ in fig. 9A, where we can see that the spike timing entropy slopes are higher than that of the spike count at all values of *n*_bins_ *≥* 1. Furthermore, the gain in entropy of a spike timing over a spike count code becomes greater as longer spike trains are considered (i.e. *n*_bins_ is increased). With the parameter values *T* = 15 ms, *τ*_ref_ = 5 ms, we plot the maximum entropy rate 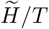 as a function of the number of neurons *N*_nrn_ in fig. 9B. For *N*_nrn_ = 10 neurons, we can see that the spike timing code achieves an entropy rate of 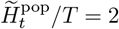 bits/ms, whereas the spike count code can only encode 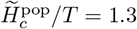 bits/ms. This analytical finding is consistent with our main computational Results, showing that spike timing codes increase the computational capacity of small populations of neurons post-synaptic to a network convergence. To reach the amount of entropy encoded by a given population employing a spike timing code, but with a spike count code, the number of neurons in the population should increase.

**Fig 9.**
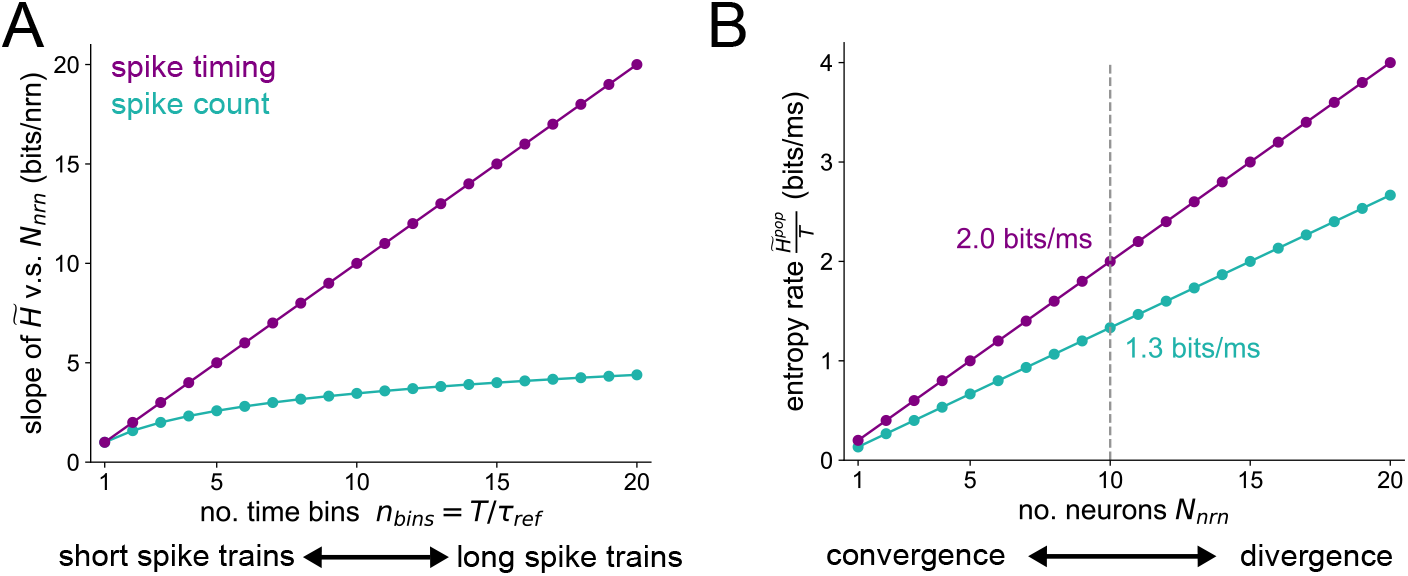
Maximum entropy of population spike codes. (A) Slope of the entropy v.s. population size curves, as a function of the number of time bins. The purple curve is simply the linear function *y* = *n*_bins_ for the spike timing code and the teal curve is the function *y* = log_2_(*n*_bins_ + 1) for the spike count code. (B) Example of the entropy rate v.s. population size for both types of spike code. We set *T* = 15 ms and *τ*_ref_ = 5 ms here so that *n*_bins_ = *T/τ*_ref_ = 3.

### Spiking neuron models

The Python package [snnTorch](https://snntorch.readthedocs.io/en/latest/readme.html) [38] was used to train and run simulations of the spiking neural networks (SNNs) studied here. The spiking neuron model that is used for all primary results is the spike response model or “alpha” neuron. We have also implemented a simple leaky integrate-and-fire neuron model, to verify that the main results are not model-dependent.

The evolution of the [alpha neuron](https://snntorch.readthedocs.io/en/latest/snn.neurons_alpha.html) is governed by the following difference equations:

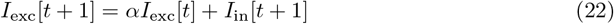

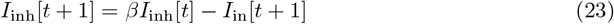

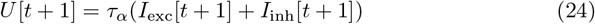

where *α* is the decay rate of the excitatory current *I*_exc_ and *β* is the decay rate of the inhibitory current *I*_inh_. The term *I*_in_ represents external current, which either comes from a stimulus or pre-synaptic spiking. The time constant for the membrane potential *U* is given by 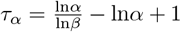. To ensure that positive inputs increase the membrane potential, we set *α > β*.

**Table 1.** Parameter values for the neurons in the alpha neuron model. The symbol *U* (*A, B*) denotes the uniform distribution between *A* and *B*.

| Alpha neuron parameter initializations |  |  |
| --- | --- | --- |
| Parameter name | symbol | value |
| Excitatory current decay rate | $\alpha$ | $\mathcal{U}(0.7, 0.9)$ |
| Inhibitory current decay rate | $\beta$ | $\alpha - 0.1$ |
| Reset membrane potential | $U_{reset}$ | 0 |
| Threshold membrane potential | $U_{thr}$ | $\mathcal{U}(0, 0.5)$ |

The [leaky integrate-and-fire (LIF) neuron](https://snntorch.readthedocs.io/en/latest/snn.neurons_leaky.html) is governed by

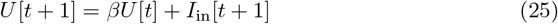

where *β* is the decay rate of the membrane potential *U* and *I*_in_ is the input current. For both the alpha and LIF neuron models, we set *U* [*t* + 1] = *U*_reset_ whenever the membrane potential reaches the spiking threshold *U* [*t*] *> U*_thr_.

**Table 2.** Parameter values for the neurons in the LIF neuron model. The symbol 𝒰 (*A, B*) denotes the uniform distribution between *A* and *B*.

| LIF neuron parameter initializations |  |  |
| --- | --- | --- |
| Parameter name | symbol | value |
| Membrane potential decay rate | $\beta$ | $\mathcal{U}(0.7, 0.9)$ |
| Reset membrane potential | $U_{\text{reset}}$ | 0 |
| Threshold membrane potential | $U_{\text{thr}}$ | $\mathcal{U}(0, 1.1)$ |

Our network’s input layer is inspired by an insect visual system with a mechanism for motion direction selectivity in two dimensions in its visual scene [69, 70]. We designed the input layer of our models to be tuned to various regions of a 2-dimensional plane (the visual scene). For the *i*^th^ neuron in the subpopulation of the input layer tuned to quadrant *q*, the input it receives is

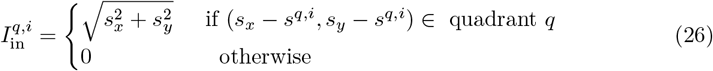

where *q* = 1, 2, 3, 4 and *i* = 1, …, *N*_in_/4. The stimulus *s* is a time-dependent vector with two components *s*_*x*_ and *s*_*y*_ denoting x- and y-positions of a moving object. The total number of neurons in the input layer is *N*_in_. We generate a set of random offsets 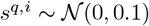 for each quadrant, independently sampled for each neuron. The purpose of this is to encourage smooth transitions between firing of the 4 subpopulations, which is more biologically-realistic than discrete switching.

### Network connectivity

To capture critical biological features, our spiking neuron model includes both excitatory and inhibitory input synapses because of their role in passing on relevant sensory information and evoking balanced motor responses in the sensorimotor pathway [20].

All layers besides the input layer of our feedforward network models solely receive inputs from neurons in pre-synaptic layers. The *i*^th^ neuron in the (*k* + 1)^th^ layer other than the input layer receives the input current

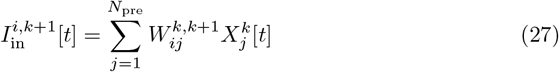

where *N*_pre_ is the number of neurons in the layer pre-synaptic to neuron *i* in layer *k, W*^*k,k*+1^ is the synaptic weight matrix from layer *k* to layer *k* + 1, and 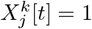 if pre-synaptic neuron *j* in layer *k* spiked at time *t* and 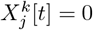 otherwise. The entries of *W*^*k,k*+1^ that are non-zero with probability *p* are distributed according to 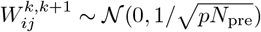. Thus, excitatory and inhibitory connections are equally probable in our model, and may both exist for the same pre-synaptic neuron and Dale’s law was disregarded for simplification. We set the connection probability to *p* = 0.7 for all models.

### Network training

The synaptic weights between layers of spiking neurons in our networks are optimized with backpropagation-thru-time (BPTT) to minimize the following loss function:

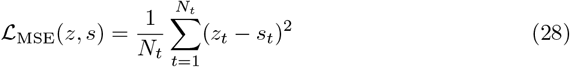

where *N*_*t*_ is the total number of time points, *s*_*t*_ is the true stimulus at time *t*, and *z* is a readout from the output layer of the form

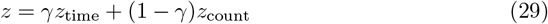

where γ = 0.5 to equally weigh spike count and timing, *z*_time_ = *W*_time_*r*_time_ and *z*_count_ = *W*_count_*r*_count_. The matrices 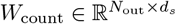 and 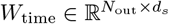 are read-out weights whose entries are initialized randomly from a normal distribution 𝒩 (0, 0.1). The symbol *d*_*s*_ denotes the dimensionality of the stimulus dynamics: either *d*_*s*_ = 1 for the 5-layer network or *d*_*s*_ = 2 for the 3-layer network. The quantities 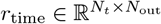 and 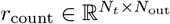 are convolutions of the output layer’s spike trains with two different kernels:

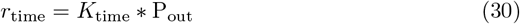

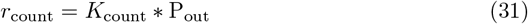

where ∗ denotes convolution. The binarized population spikes of the output layer 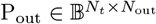 (where 𝔹 = {0, 1}) are convolved with the kernels *K*_time_ and *K*_count_, which are of the form

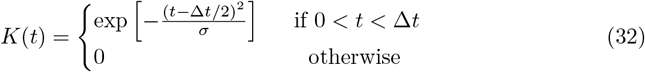

where Δ*t* = 10 ms for 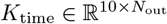 and Δ*t* = 70 ms for 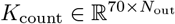. We chose Δ*t* = 10 ms as the scale of the timing convolution since only 1-3 spikes typically fall within this window, and smaller Δ*t*’s resulting in poor training. The value Δ*t* = 70 ms was chosen for the count convolution since multiple spikes usually fall within this window. Values larger than Δ*t* = 70 ms for the count convolution resulted in poorer training. The standard deviation is set to *σ* = 0.1 ms^2^ for both kernels.

The read-out weights *W*_time_ and *W*_count_, as well as the membrane decay rates *α* and *β* and synaptic weights (see Spiking neuron models) of the spiking neural network, are trained during back-propagation to minimize the mean-squared error. A plot of the MSE loss over training is shown in fig. 10, as well as an example of the read-out compared to the stimulus after training.

**Fig 10.**
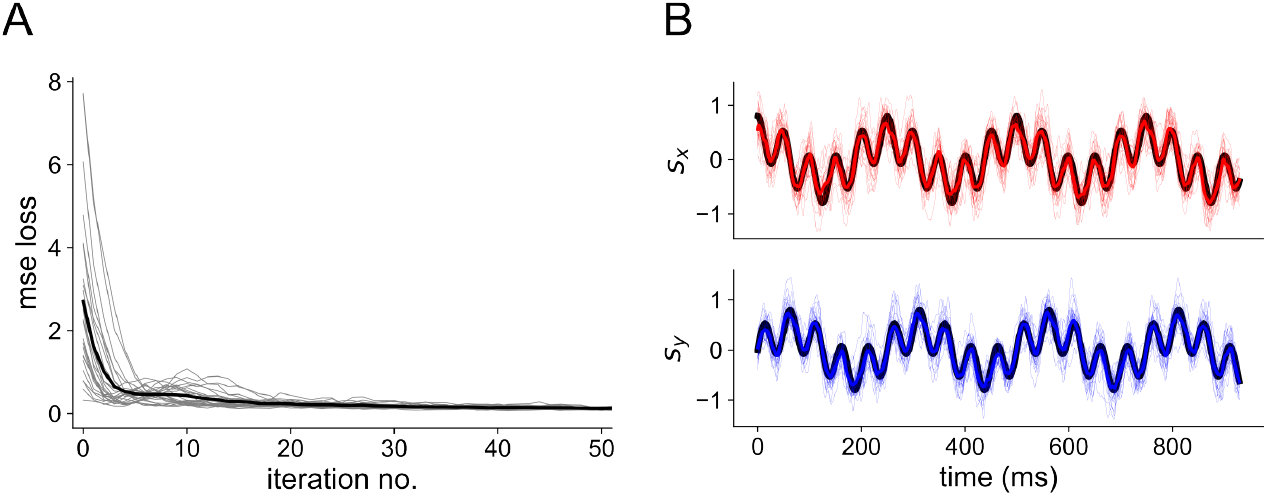
Network training. (A) Reduction of MSE loss through training with BPTT. Thin gray traces show individual network seeds, thick black trace shows the average across all 25 seeds. (B) Readout after training 3-layer networks with *N*_in_ = *N*_h_ = *N*_out_ = 100 to the 4 Hz + 20 Hz sum of sines stimulus. Colored traces are for the readout; the black trace denotes the true stimulus presented to the network. The top shows the *x*-dimension of the stimulus and the bottom shows the *y*-dimension.

### Decoding analysis

In order to determine how the population responses of layers in our network model relate to stimuli, we trained and tested a decoder [42]. In particular, long short-term memory (LSTM) and gated recurrent unit (GRU) networks were used to predict the stimulus at time *t* based on the neural response during time [*t, t* + *T*] binned at resolution Δ*t*. In other words, the stimulus value at the beginning of the spike train is the value we use the spike train to decode. To further clarify this process, suppose a neural recording of *t*_*f*_ = 10 time steps results in the following spike train:

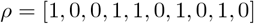

where “0” represents no spike and “1” represents spike. Sliding a rectangular window of width *T* = 8 over this spike train results in

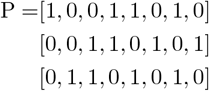

Each of these are then sub-divided into bins of size Δ*t*. If Δ*t* = *T* = 8, the binned response *R* becomes a vector of spike counts over the response window *T* :

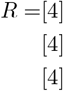

If Δ*t* = 4, then the binned response is

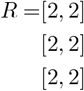

If Δ*t* = 2, then the binned response is

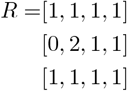

And if Δ*t* = 1, then the binned response becomes identical to the original binary spike train P.

The above matrix *R* has size (*n*_samples_ × *n*_bins_) where *n*_samples_ = *t*_*f*_ *− T* + 1 = 10 *−* 8 + 1 = 3 and *n*_bins_ = *T/*Δ*t*. If instead of 1 neuron, we have recordings from *N*_nrn_ neurons (as in our population decoding analyses), the same procedure is performed on each neuron’s spike train and their resulting matrices are stacked together to form a tensor of dimension (*n*_samples_ × *n*_features_ × *n*_bins_) where *n*_features_ = *N*_nrn_. This tensor is used to decode the stimulus over time. The dimension along which spikes are binned at resolution Δ*t* is treated as a hidden state for the LSTM and GRU decoders, so that decoding depends on specific spike sequences. The stimulus is stored as a matrix *S* of size (*n*_samples_ × *d*_*s*_) where *d*_*s*_ is the dimension of the stimulus, either 1 or 2 here. The task of decoding is to find a function *f* that forms an estimate *Ŝ* = *f* (*R*) of the true stimulus *S*, minimizing the error ∑_*i,j*_ (*ŝ*_*ij*_ *− s*_*ij*_)^2^. In our analysis, the control parameter Δ*t* is varied to modulate the time resolution with which spikes are counted. When Δ*t* = 1, there is no difference between P and *R*, and the specific timing of every spike is preserved. As Δ*t* is increased, spike timings within the larger window of size *T* become increasingly blurred. The maximum value Δ*t* = *T* results in a vector *R* where each entry is the number of spikes that occurred in the respective time window of duration *T*. On the other hand, as Δ*t* decreases, the code becomes more dependent on spike timing than spike count.

We used the Python package [keras](https://github.com/keras-team/keras) to perform the decoding with the LSTM and GRU networks. Cross-validation was performed by maximizing the validation accuracy using Bayesian optimization [71] to select hyperparameters.

### Single-neuron information theoretic analysis

We follow Putney et al. [25] for the single neuron mutual information analysis. Briefly, the idea is to compute the mutual information *I*_*m*_ between motor output *m* and single-neuron response *R* via:

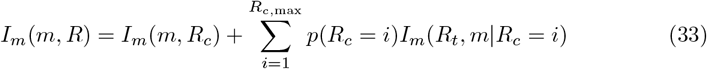

where *R*_*c*_ is the spike count, *R*_*t*_ is the spike timings, and *m* is the first two principal components of the motor output (forces/torques generated by the wing muscles during hover feeding). The first term in eq. (33) is what we label the “spike count” information and the second term is the “spike timing” information in the single-neuron analyses of the 5-layer network results. In our implementation, 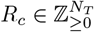 where ℤ_*≥*0_ denotes the set of non-negative integers and *N*_*T*_ = (*N*_*t*_*/T*) is the number of non-overlapping response windows of duration *T* falling within the experiment or simulation of duration *N*_*t*_. For the moth experiments, *T* = 50 ms is the same as the wingstroke period of the animal, so *N*_*T*_ equals the number of wingstrokes in this context. The spike timing matrix 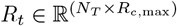 contains the spike timings within each response window where *R*_*c*,max_ is the maximum number of spikes observed in a single wing stroke. The quantity *p*(*R*_*c*_ = *i*) denotes the probability that a spike count of *i* was observed. The mutual information in spike count *I*_*m*_(*m*; *R*_*c*_) and the mutual information in spike timing, conditioned on spike count, *I*_*m*_(*R*_*t*_, *m*|*R*_*c*_ = *i*) were both estimated numerically using the Kraskov-Stögbauer-Grassberger (KSG) method [72] (see Assocation measures). Since there is no “motor output” for our network model, we performed this analysis by substituting the stimulus *s* for the motor output *m* in eq. (33). As the stimulus represents the position of a target that moths follow, it is reasonable to assume that the stimulus information is directly reflected in the motor output.

### Assocation measures

To quantify the amount of information between stimulus and response in our population decoding analyses, we employ various association measures between the true stimulus *S* and decoded stimulus *Ŝ* = *f* (*R*) based on the response *R*. If we define *I*_*m*_(*X, Y*) as the mutual information between random variables *X* and *Y*, the data-processing inequality states that *I*_*m*_(*S, Ŝ*) ≤ *I*_*m*_(*S, R*) since *Ŝ* cannot gain information about *R* [73]. For large populations of neurons and small Δ*t*’s, the response matrix *R* becomes very high dimensional, rendering the quantity *I*_*m*_(*S, R*) difficult to estimate directly [74]. Thus, we instead estimate the quantity *I*_*m*_(*S, Ŝ*) which forms a lower bound on the true mutual information of interest *I*_*m*_(*S, R*). This is done via the Kraskov-Stögbauer-Grassberger (KSG) method [72], employed via scikit-learn. For the single-neuron mutual information calculations, we used the Julia package Associations.jl. In addition to mutual information, which is a nonlinear measure of association between variables, we also show results with the coefficient of determination *R*^2^ (decoding accuracy) which is a linear association measure.

## Supporting Information

**S1 Fig. Version of fig. 2 but with GRU decoder**

**S2 Fig. Version of fig. 2 but with LIF spiking model**

**S3 Fig. Version of fig. 3 but with mutual information**.

**S4 Fig. Version of fig. 3 but with GRU decoder**.

**S5 Fig. Version of fig. 3 but with GRU decoder and mutual information**.

**S6 Fig. Version of fig. 3 but with LIF neuron model**.

**S7 Fig. Decoding analysis of 5 layer model receiving 1 Hz stimulus**

**S8 Fig. Single-neuron information across layer of the 5-layer model for three values of** *k*, **the number of nearest neighbors in the KSG method**.

**S9 Fig. Single-neuron information across layer of the 5-layer model for various data set sizes** *n*.

**S10 Fig. Robustness of the single-neuron information estimates**

**S11 Fig. Interaction info in experimental data v.s. 5-layer model**

## Acknowledgments

We thank Soon Ho Kim for insightful comments and discussions on the manuscript.

## Supporting Information

### Generalizability of the three-layer results across spike generators and decoders

#### Output layer result robustness

This section contains results verifying that our result in fig. 2 is robust to a different spiking (LIF) model and decoder (the gated recurrent unit, GRU).

**Fig S1.**
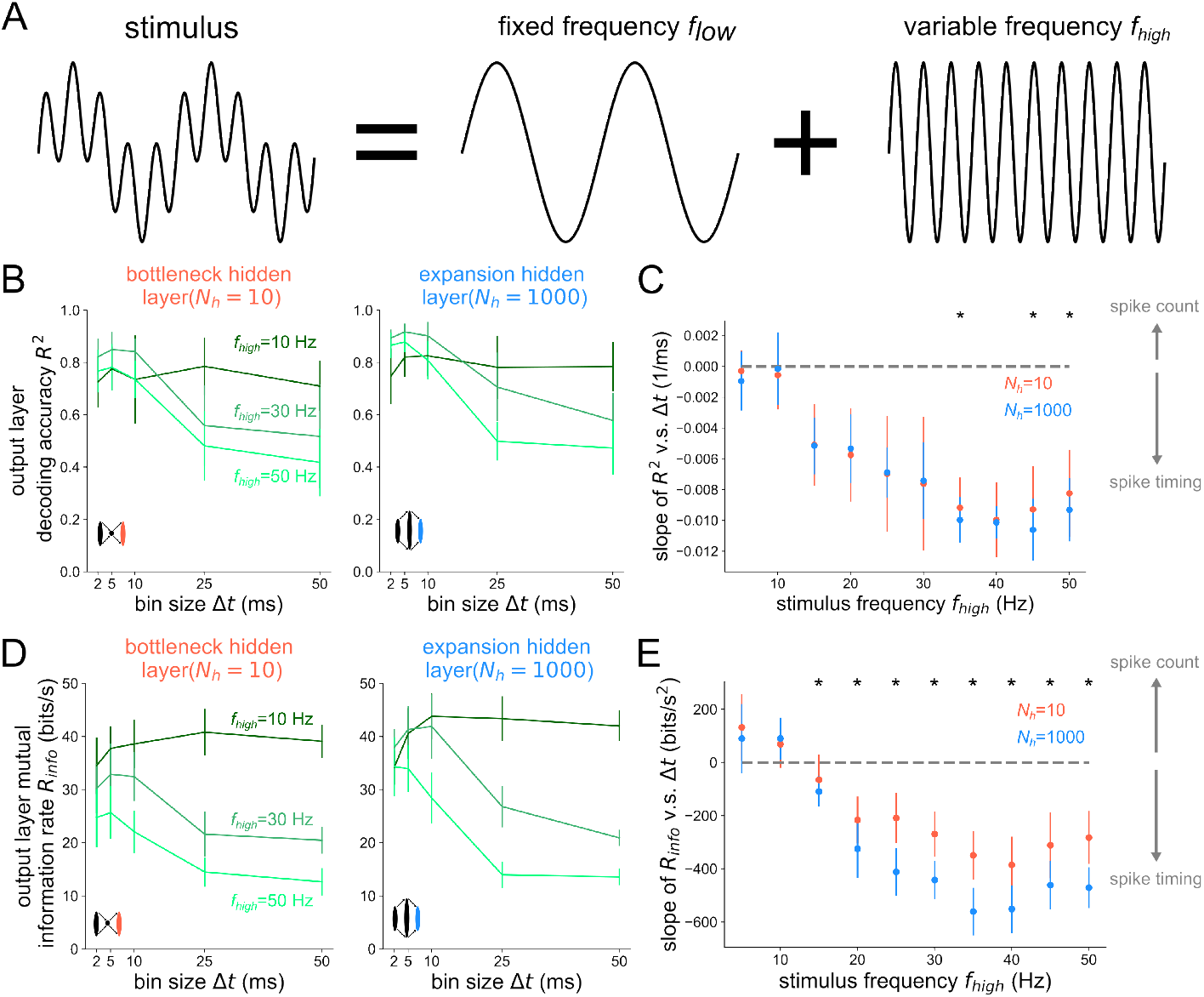
Version of fig. 2 but with GRU decoder.

**Fig S2.**
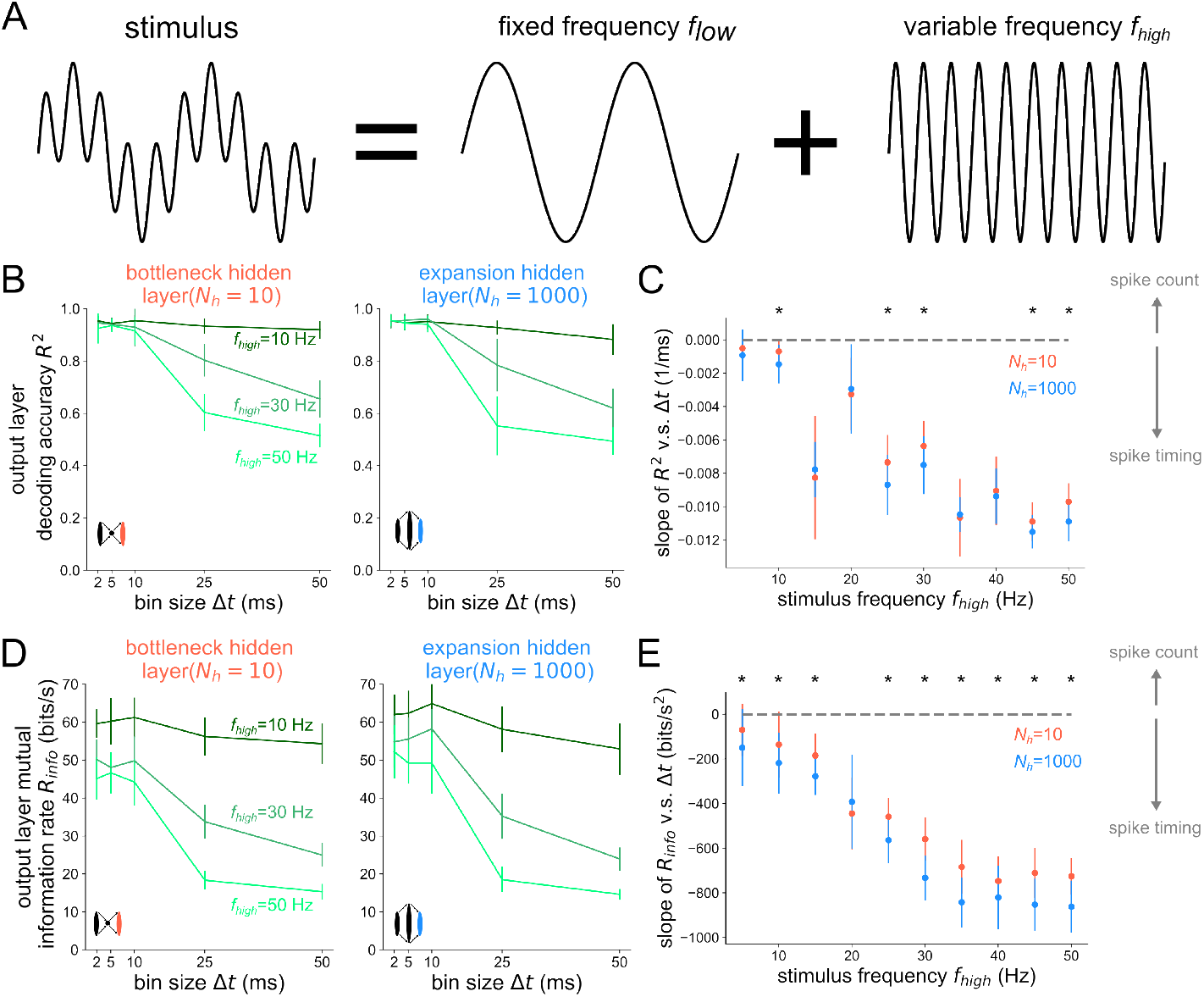
Version of fig. 2 but with LIF spiking model.

#### Hidden layer result robustness

This section contains results that verify the result in fig. 3 is robust to a different spiking model, decoder, and association metric.

**Fig S3.**
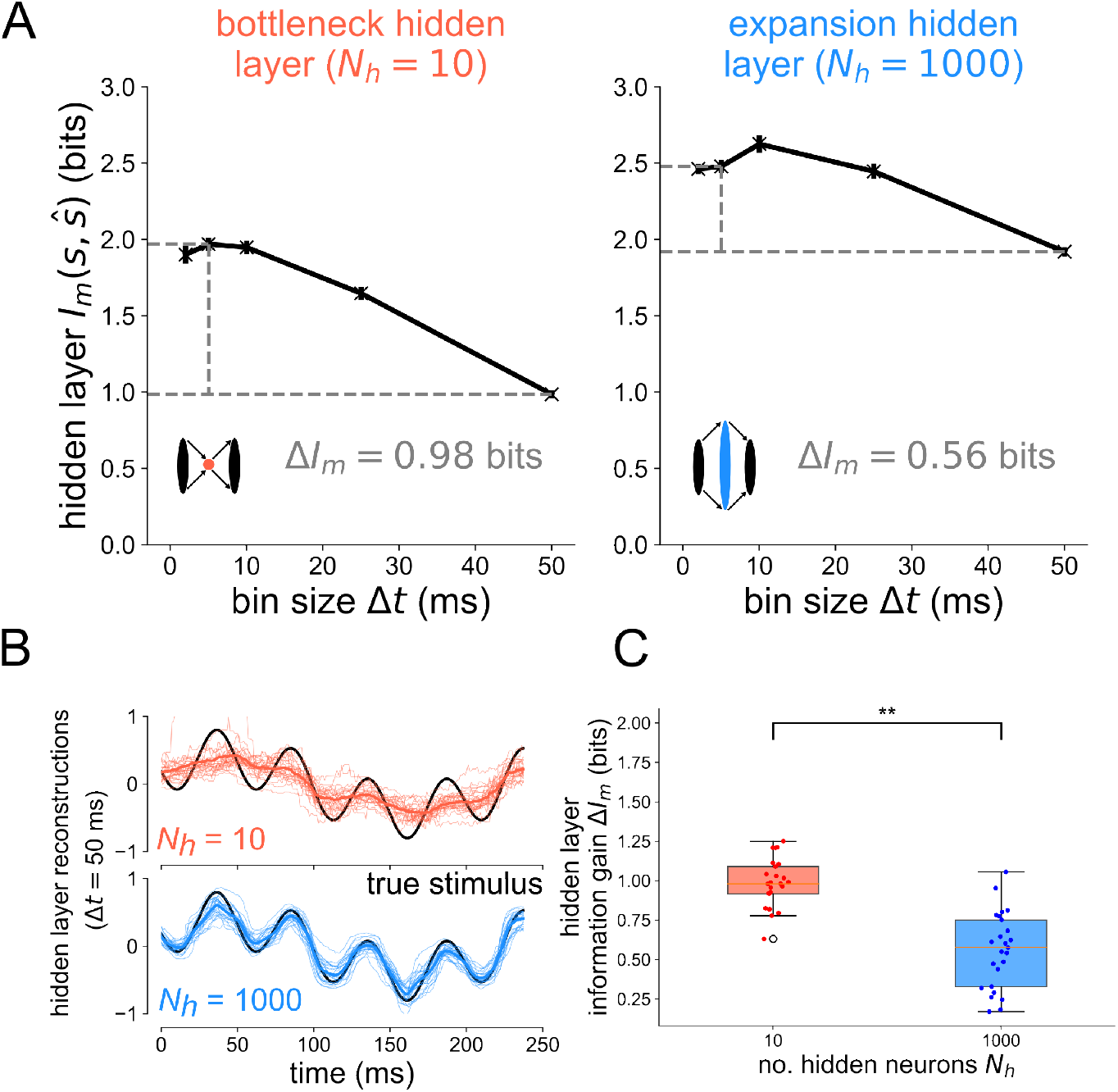
Version of fig. 3 but with mutual information.

**Fig S4.**
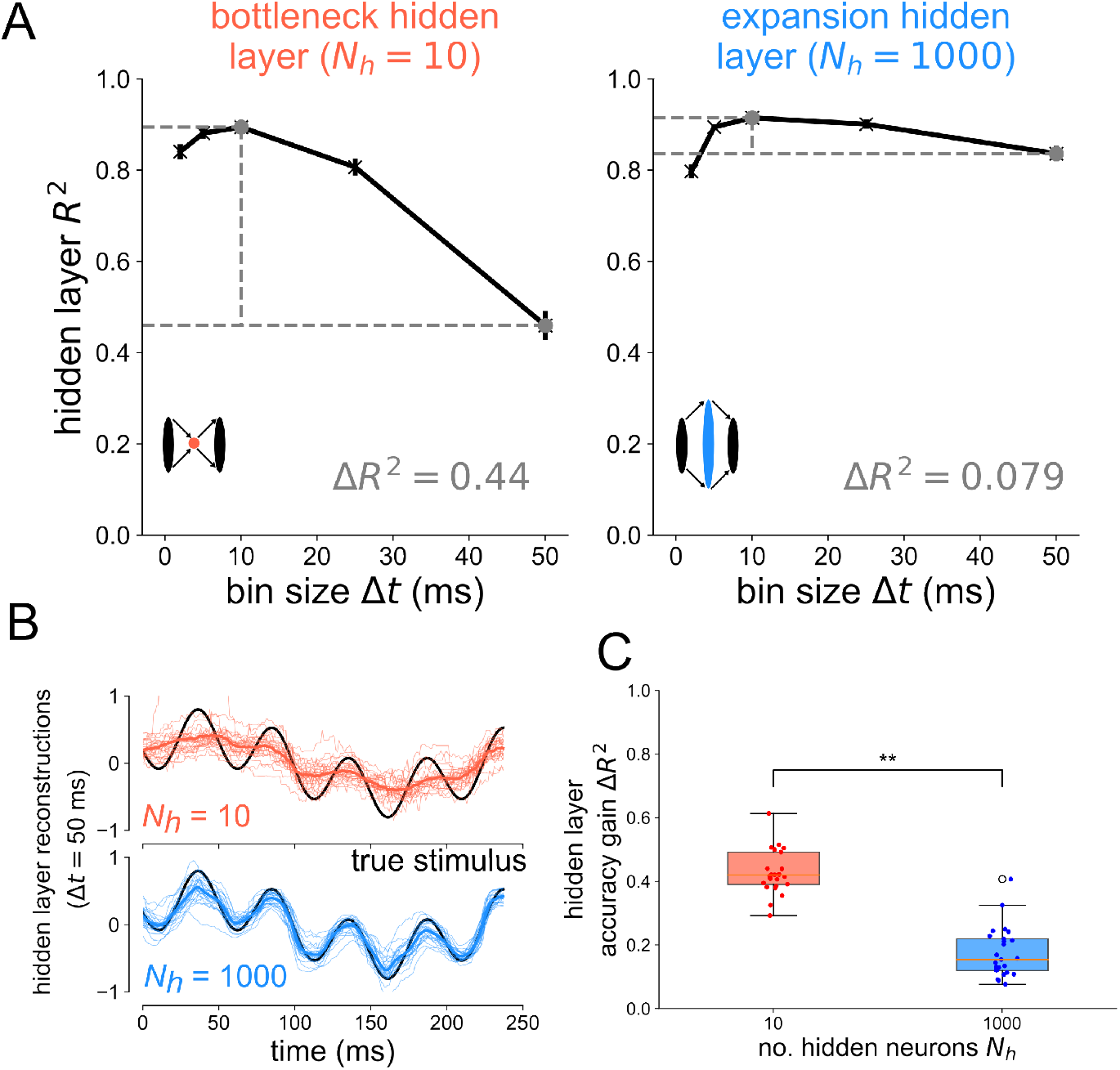
Version of fig. 3 but with GRU decoder.

**Fig S5.**
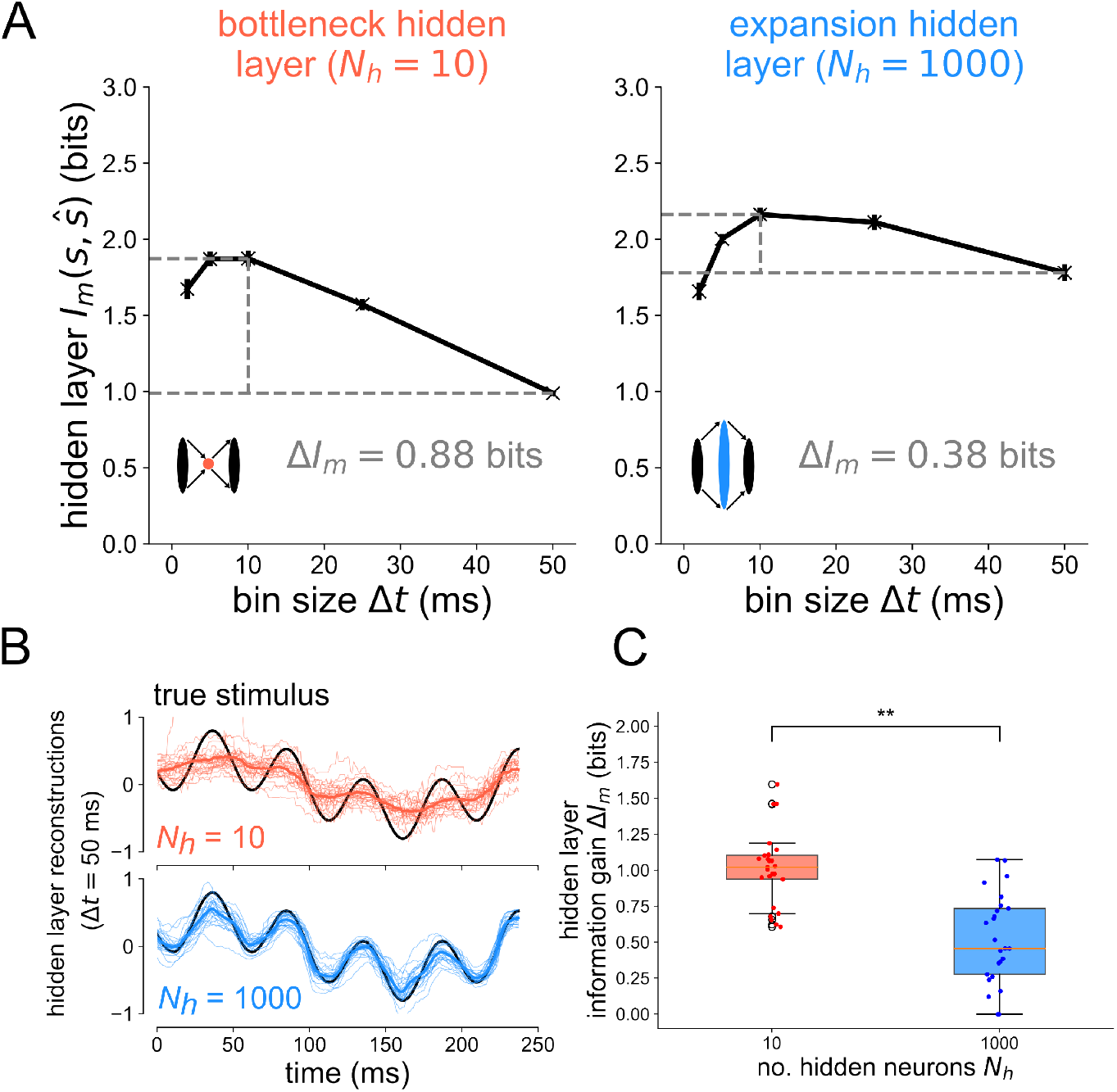
Version of fig. 3 but with GRU decoder and mutual information.

**Fig S6.**
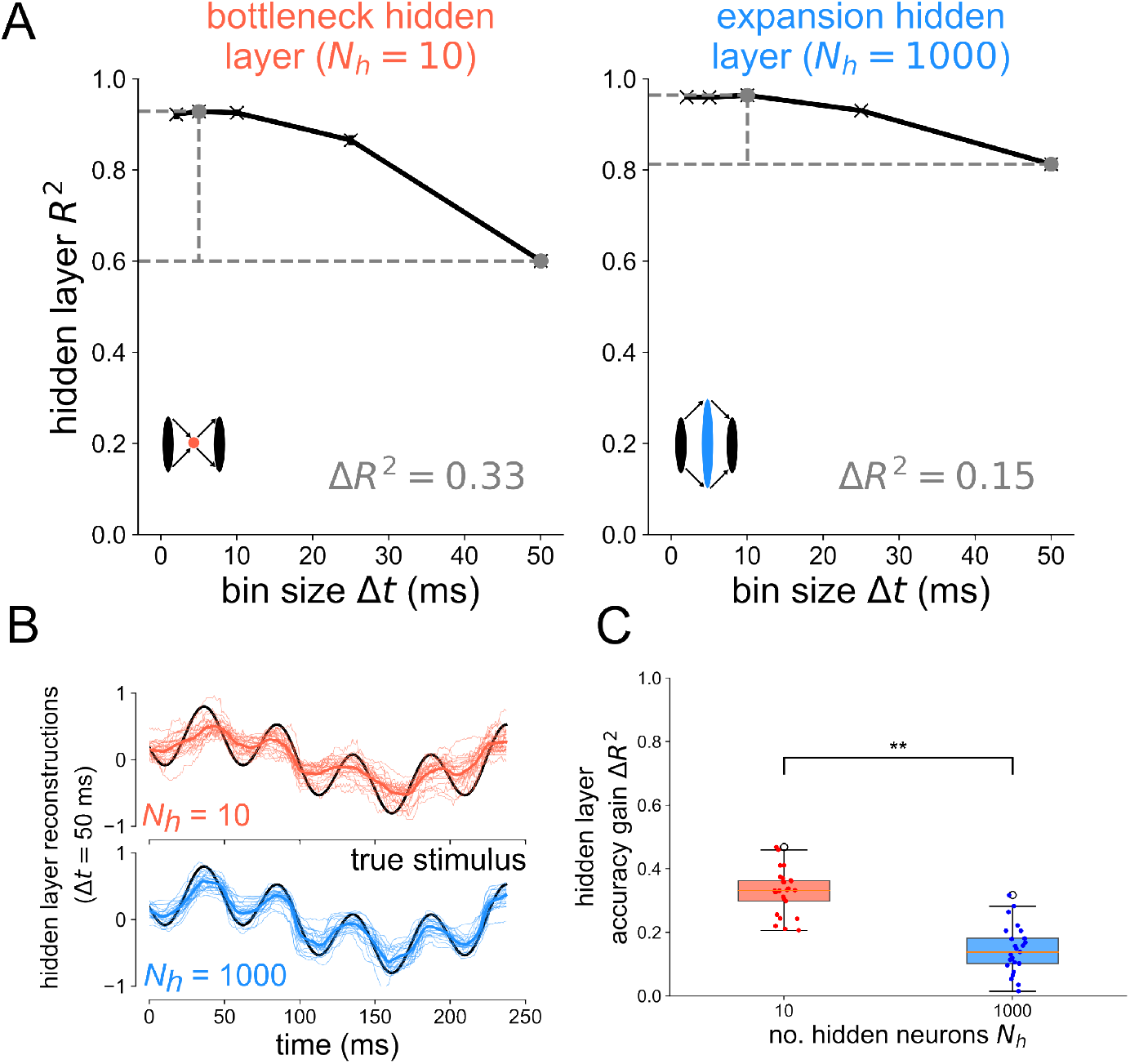
Version of fig. 3 but with LIF neuron model.

#### Five-layer model 1 Hz stimulus decoding analysis

This section shows the result of the population decoding analysis from the 5-layer network during a 1 Hz sinusoidal stimulus.

**Fig S7.**
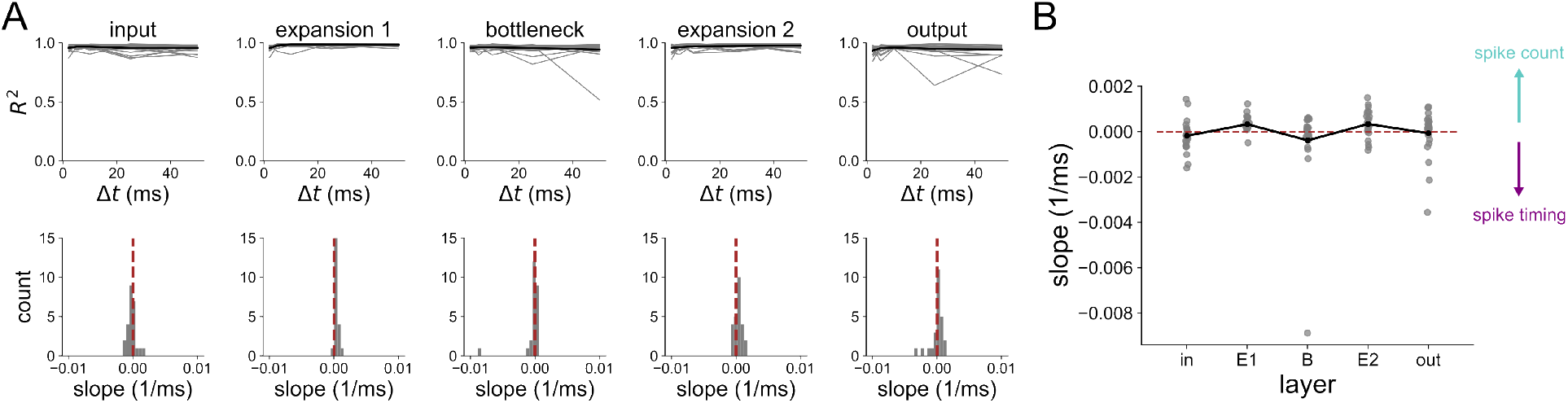
Decoding analysis of 5 layer model receiving 1 Hz stimulus. (A) Decoding accuracy *R*^2^ between true stimulus *s* and estimated stimulus *ŝ* from spikes binned at resolution Δ*t* (top) in each layer of the 5-layer model. Each gray line shows the results from one of 25 network seeds; the black curve is the mean. Slopes of the *R*^2^ v.s. Δ*t* curves in each layer (bottom). Vertical line at slope=0 is plotted for reference. (B) Slope distributions versus layer, gray dots are individual network seeds. The black lines connect the means at each layer.

#### Single-neuron mutual info robustness

This section contains results verifying that the single-neuron mutual information estimates in 5-layer network are robust to variation in the hyperparameter and dataset. In figure S8 below, we show that the single-neuron mutual information estimates across layer are robust to a range of *k*, the number of nearest neighbors hyperparameter of the KSG method.

**Fig S8.**
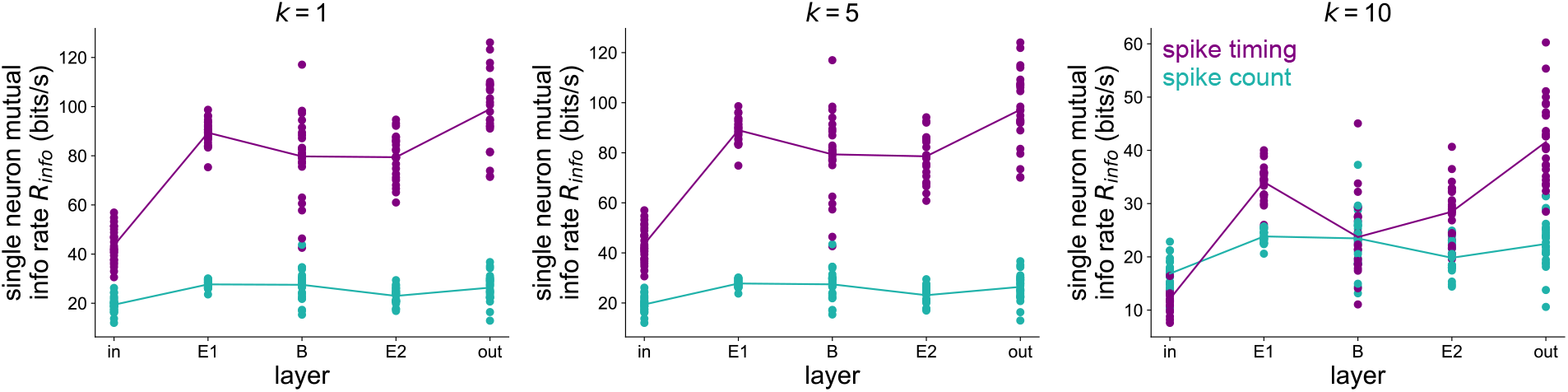
Single-neuron information across layer of the 5-layer model for three values of *k*, the number of nearest neighbors in the KSG method. Each dot represents the outcome of the single-neuron mutual information analysis for a single 5-layer network seed, pooled across all neurons in the output layer. 25 network seeds used here.

Figure S9 shows how the single-neuron mutual information estimates vary with the number of data points used from the output layer of the hawkmoth visuomotor model trained to a 1 Hz sinusoidal stimulus. The general feature where spike count information is similar to spike timing information in the input layer but much lower than spike timing information in the output layer is preserved across data set sizes.

**Fig S9.**
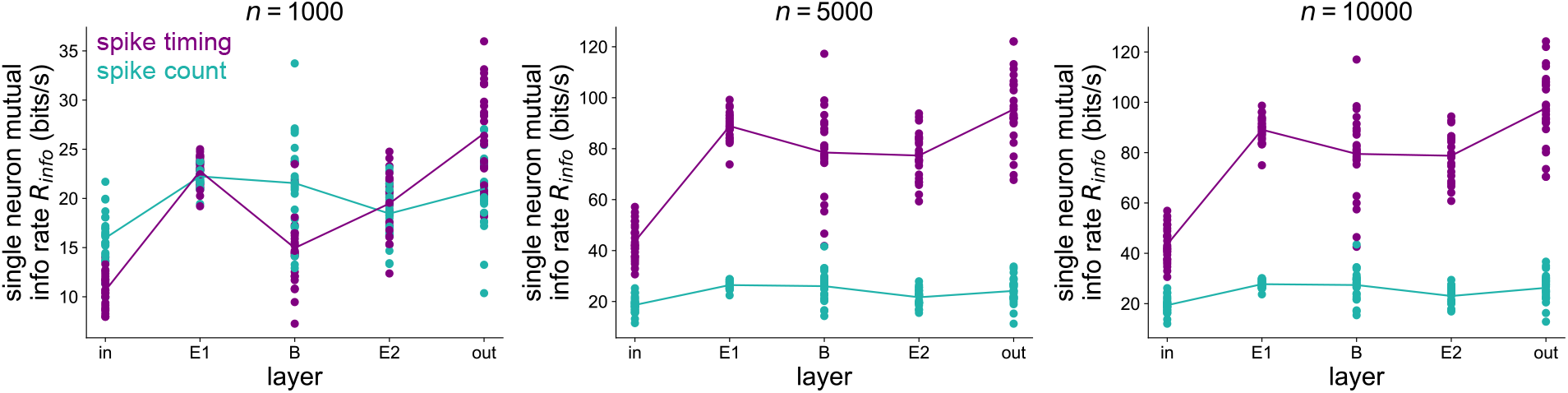
Single-neuron information across layer of the 5-layer model for various data set sizes *n*. Each dot represents the outcome of the single-neuron mutual information analysis for a single 5-layer network seed, pooled across all neurons in the output layer. 25 network seeds used here.

The single-neuron information analysis of the output layer is shown against *k* and *n* in figure S10. Over the ranges of *k* and *n* tested, the spike timing information is always significantly higher than the spike count information. In the main results, we use *k* = 3 and *n* = 10 *×* 10^3^.

**Fig S10.**
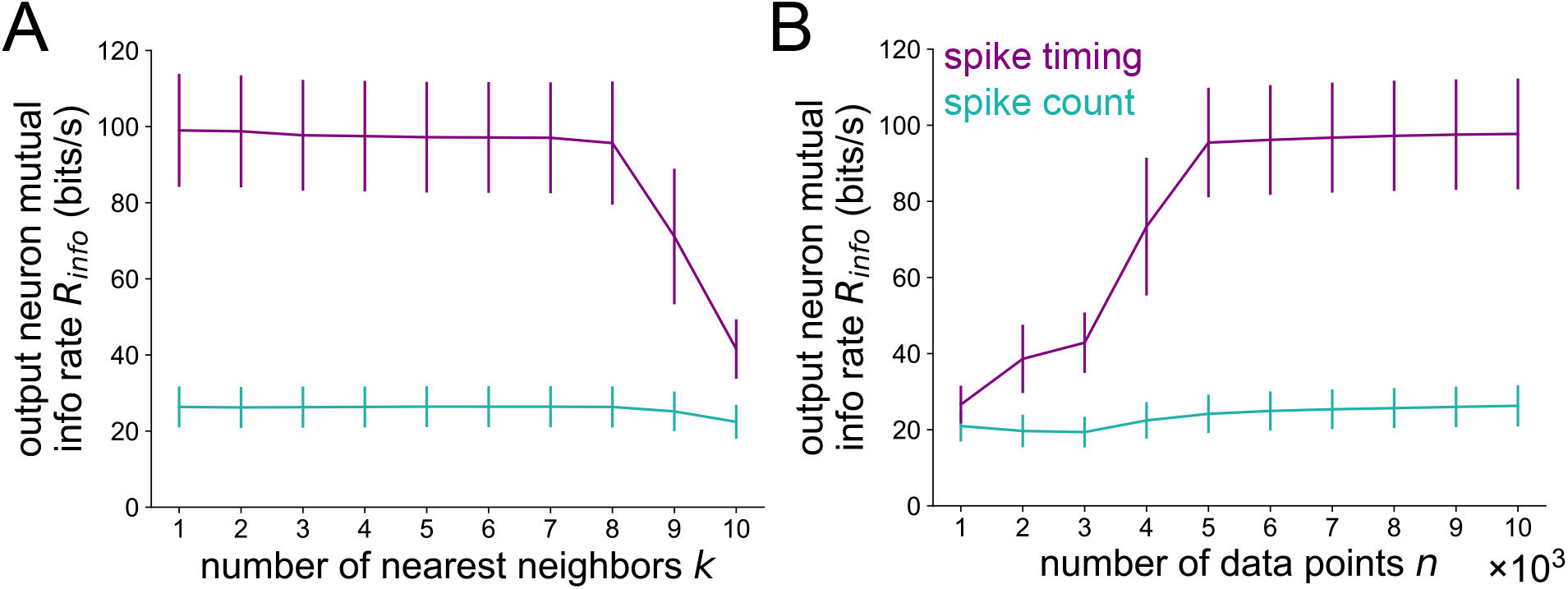
Robustness of the output layer results. (A) across number of nearest neighbors *k* and (B) across data set size *n*. Error bars represent standard deviations over 25 independent simulations.

#### Interaction information in hawkmoth model

Another key finding from the previous analysis of Putney et al [25] was that most of the redundancy in the mutual information that moth muscles share with the motor output was in spike timing, not spike count. The authors quantified this through the interaction information between pairs of neurons *A* and *B* with motor output *m*, as defined by

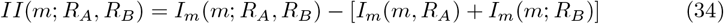

where *I*_*m*_(*m*; *R*_*A*_) is the single-neuron mutual information as defined by equation 33 and *I*_*m*_(*m*; *R*_*A*_, *R*_*B*_) is the joint mutual information that the responses of *A* and *B* (*R*_*A*_ and *R*_*B*_) share with the motor output *m*, as defined by

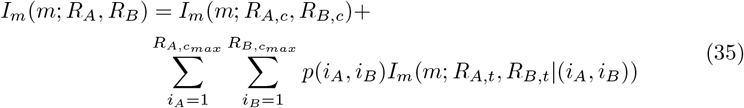

where *R*_*A,c*_ and *R*_*B,c*_ are the spike counts of neurons *A* and *B*, respectively. The maximum spike counts of neurons *A* and *B* are 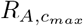 and 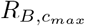, respectively. The responses *R*_*A,t*_ and *R*_*B,t*_ are the spike timings of these neurons, which are conditioned on the joint spike count probability distribution *p*(*i*_*A*_, *i*_*B*_). Negative interaction information indicates net redundancy between the pair of neurons. Positive interaction information indicates net synergy. For more details, see ref. [25]. Following Putney et al, we performed this analysis and partitioned the interaction information into spike count and spike timing contributions. We show the results from the original experimental data in the top row of Fig. S11. In the bottom row, we show the result of this analysis performed on the output layer of our 5-layer convergent/divergent network model trained to 1 Hz stimulus. As before, since there is no “motor output” for the model, we use the stimulus *s* instead of the motor output *m* in equations 34 and 35, assuming that the motor output reconstructs the stimulus position during tracking. In both the moth data and model data, a majority of the pairwise redundancy is in spike timing instead of spike count, as demonstrated by large negative *II*_*time*_ and *II*_*count*_ *≈* 0 in Fig. S11.

**Fig S11.**
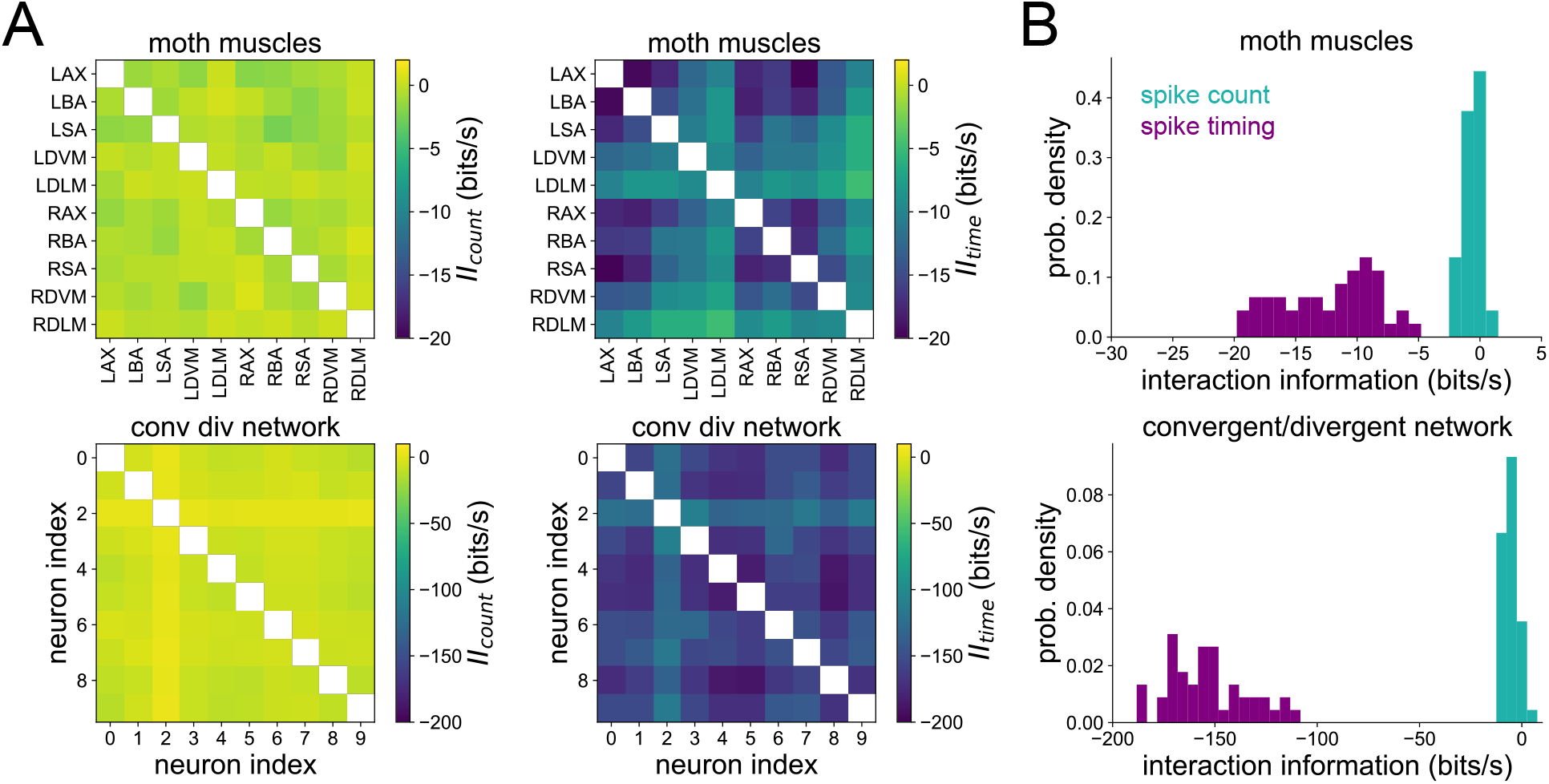
Interaction info in experimental data v.s. 5-layer model. (A) Interaction info from spike count (*II*_*count*_) and spike timing (*II*_*time*_) in experimental data (top) and the output layer of our convergent-divergent 5-layer model of the hawkmoth visuomotor pathway (bottom). (B) Interaction info in spike count and timing pooled across all moth muscles (top) and output layer neurons in the model (bottom).

